# Reconstructing stochastic cell population trajectories reveals regulators and heterogeneity of endothelial flow-migration coupling driving vascular remodelling

**DOI:** 10.1101/2023.05.30.542799

**Authors:** Wolfgang Giese, André Rosa, Elisabeth Baumann, Olya Oppenheim, Emir B. Akmeric, Santiago Andrade, Irene Hollfinger, Eireen Bartels-Klein, Silvanus Alt, Holger Gerhardt

**Affiliations:** Max Delbrück Center for Molecular Medicine in the Helmholtz Association, Berlin, Germany; DZHK (German Center for Cardiovascular Research), Berlin, Germany; Molecular Medicine and Gene Therapy, Lund Stem Cell Centre, Lund University, Lund, Sweden; Charité - Universitätsmedizin Berlin, Berlin, Germany

**Keywords:** vascular remodelling, endothelial cells, Cdc42, collective migration, flow-migration coupling, migration dynamics

## Abstract

Emerging concepts of developmental vascular remodelling recently identified that selectively labelled sprouting tip cells and/or venous endothelial cells (ECs) accumulate in developing arteries, suggesting directional migration of specific ECs drives artery formation. However, a general population analysis and detailed quantitative investigation of migratory mechanisms is so far lacking. Here, we developed a universally applicable quantitative approach and a computational model allowing to track and simulate stochastically labelled EC populations irrespective of labelling density and origin. Dynamic mapping of EC distributions in a bespoke coordinate system revealed how ECs move during the most active remodelling phases in the mouse retina. Simulation and parameter sensitivity analysis illustrated that the population shift from veins to arteries cannot be explained by random walk, but best fits to a tuneable dual force-field between shear-force induced directionality against blood flow, and hypoxia mediated VEGFA-gradients. High migration rates require only weak flow-migration coupling, whereas low migration rates require strong coupling to the flow direction. Functional analysis identified Cdc42 as the critical mediator of overall population movement from veins to arteries, yet with surprising heterogeneity suggesting the existence of distinct cell populations. This new quantitative understanding will enable future tailored intervention and tuning of the remodelling process.

## Introduction

The mechanisms establishing the hierarchical patterning of arteries, arterioles, capillaries, venules and veins to effectively irrigate all tissues appear highly complex and remain incompletely understood. Large arteries co-align with major nerve trajectories through joint local tissue guidance cues (Mukouyama et al., 2002), and repulsive cues appear to segregate venous from arterial cells (Eichmann et al., 2005). Many signalling molecules and transcription factors that influence arterio-venous fate decisions have been identified, but exactly how endothelial cells (ECs) coordinate their collective behaviour during the remodelling process that establishes the mature vessel networks has remained obscure.

Recent lineage tracing studies in the mouse retina identified migration of tip cells to arteries, and also direct migration of cells from veins to arteries along existing vessel connections through the remodelling vessel plexus (Lee et al., 2022). Unlike tip cell migration towards hypoxic tissue environments along VEGFA gradients, the migratory behaviour from the tip position or the vein towards arteries occurs in the opposite direction, towards higher oxygen tension and thus away from higher VEGFA concentrations. Recent work by the lab of Claudio Franco demonstrated that these opposing directional attraction forces appear to compete, involving force transmission through cell-cell junctions and basal adhesions (Barbacena et al., 2022).

Studies in both retina and zebrafish, as well as mouse coronary arterial development indicate that the directional movement of cells towards arteries is in part controlled by Cxcl12 expression in arterial cells attracting Cxcr4 expressing pre-arterial cells in the plexus or from the tip region (Ara et al., 2005; Ivins et al., 2015). An additional, or alternative attractive cue is provided by directional blood flow. ECs polarize and migrate against the direction of blood flow up the gradient of increasing shear levels (Vion et al., 2021). Yet, whether all ECs follow the same principle, and how shear induces this migratory behaviour is poorly understood. Recent studies highlight the importance of junctional dynamics to enable these intercalating cell movements, and identified the actin nucleation factor Wasp to be at least in part responsible for cells to move from veins to arteries in zebrafish (Rosa et al., 2022).

Unlike zebrafish embryos, mammalian tissue development is challenging to image over time. Blood vessel development in three dimensional networks occurs at depths that are difficult to reach with light microscopy, and alternative whole tissue or body imaging approaches still lack resolution to depict cells and microvascular networks (Ueda et al., 2020). Development and remodelling of vessels also occur over days and weeks, making it very difficult to follow individual cells over time (Rust et al., 2019). Labelling cells at a given time point with genetic tools allows to trace cells from their point of labelling to any future location even if continuous imaging is not possible. The ability to label cells at precise locations, however, depends on the availability of selective genetic tools, i.e. promoters/enhancers, that can drive the expression of inducible recombinases in a discrete cell population at a precise time point. Recent work by the Simons lab generated the first vein specific inducible Cre-line, illustrating migration of venous ECs both during developmental remodelling and after pathological reactivation (Lee et al., 2021). However, quantitative information is so far lacking.

To gain quantitative insight into the dynamics of EC population movements during vascular remodelling and artery-vein formation, we developed a general coordinate system and created spatial maps of EC distributions over time. This analytical approach can be used even with non-selective stochastic recombination events to trace spatiotemporal population movements. We utilized inducible Cre-lox recombination of fluorescence reporter transgenes under the control of the Vegfr3 promoter to label scattered ECs in the developing plexus and veins, but not arteries (Martinez-Corral et al., 2016). Through detailed analysis of a time-series after label induction, we identified changes in EC localisation over time and space. Our analysis identified an EC population shift from veins to arteries and highlighted an emerging population that enters the artery. Computational simulation of stochastically labelled ECs randomly seeded in a vein-plexus-artery sector of a segmented retina illustrated that the observed population shift cannot be explained by random walk, no matter how motile ECs are. Only the combination of a dual force-field comprising a simulated hypoxia patterned VEGFA gradient, and the flow-mediated shear stress gradient lead to a close match between simulation and experimental observation. Phase diagram analysis of the movement rate versus the coupling strength to the two force fields predicted that already a low coupling strength is sufficient to drive ECs towards the artery if the movement rate is sufficiently high. In the case where ECs can only migrate very slowly, the model predicts that a high coupling rate, meaning a strong directional bias against blood flow, is required to reach the artery.

To test the model predictions, we experimentally investigated the role of two key small Rho GTPases, Cdc42 and Rac1, in the context of this migratory behaviour. Therefore, we conditionally inactivated Rac1 or Cdc42 in stochastically labelled ECs *in vivo*, using the same Cre-lox system under Vegfr3 control, in conjunction with the floxed alleles of Rac1 and Cdc42 (Chrostek et al., 2006; Wu et al., 2006). Rac1 depletion caused only a minor delay in migration, and a surprising late acceleration to reach the artery. Cdc42, however, appeared uniquely essential for cells to reach the artery, with no evidence for any directional movement against the shear gradient. Even at very high recombination rates, when most ECs in the retina were labelled and thus Cdc42 depleted, the arteries remained devoid of labelled cells. *In vitro* experiments confirmed these findings and illustrated that Rac1 depletion led to slower migration both in a scratch wound assay as well as in a flow assay, yet with retained directionality. Cdc42 depletion in contrast not only led to even stronger migration defects, but completely abrogated the ability of ECs to migrate directionally under flow. Thus, although classically Rac1 is known as the master driver of lamellipodia driven migration processes, and Cdc42 more involved in filopodia-driven processes, endothelial flow-migration coupling in vascular remodelling is exquisitely sensitive to Cdc42 function, without which, ECs cannot collectively move against flow. Understanding the quantitative and qualitative mechanisms and players regulating endothelial population movements is crucial to unravel pathomechanisms in vascular malformations and identify targets for selective interventions.

## Results

### Stochastically labelled endothelial cells move directionally towards arteries in the developing mouse retina

Vascular development and remodelling in mammalian tissues occurs over days to weeks. Tracking migratory patterns of labelled individual ECs over time is a powerful tool for uncovering the principles of dynamic EC rearrangements, but usually requires live imaging or at least the ability to image the same tissue repeatedly. To study the migratory behaviour of ECs during vascular remodelling in fixed tissues, such as the mouse retina model, where each individual retina can only be imaged at a single developmental time point, we employed genetic recombination-based lineage tracing combined with computational cell population analysis. We took advantage of a transgenic mouse model expressing Cre recombinase (Cre) fused to a mutant estrogen ligand-binding domain (ERT2) (Feil et al., 1996) that requires the presence of tamoxifen for activity under the control of the Vegfr3 promoter Vegfr3Cre^ERT2^ (Martinez-Corral et al., 2016; Tammela et al., 2011). Vegfr3 is strongly expressed in the lymphatic endothelium of adult mice, but also more widely in activated vascular ECs during embryonic and early postnatal development. In the mouse retina, which lacks lymphatic vessels (Chen, 2009; Tammela and Alitalo, 2010), Vegfr3 is expressed in endothelial tip cells and ECs in the developing vein, as well as scattered ECs throughout the primitive vascular plexus. ECs in arteries, however, do not express Vegfr3. Accordingly, acute Cre-mediated recombination of the ubiquitous Cre-reporter mTmG (Muzumdar et al., 2007) after a single injection of tamoxifen turned off membrane-tethered Tomato expression, and turned on mGFP expression in ECs in the vein, tip cells and plexus. Similar specificity of localized activation can be observed after acute recombination at later stages, as long as vascular development is actively ongoing (Figure S1, P8-P9). Recombination in arteries could not be observed acutely after tamoxifen injection at any time point, confirming that Vegfr3 is not actively expressed in ECs in arteries. Intriguingly, however, retinas dissected up to 4 days after tamoxifen injection consistently show GFP-positive ECs in arteries, suggesting that recombined ECs migrate from the above-mentioned locations to arteries in the remodelling retinal vasculature (Figure 1A).

**Figure 1:**
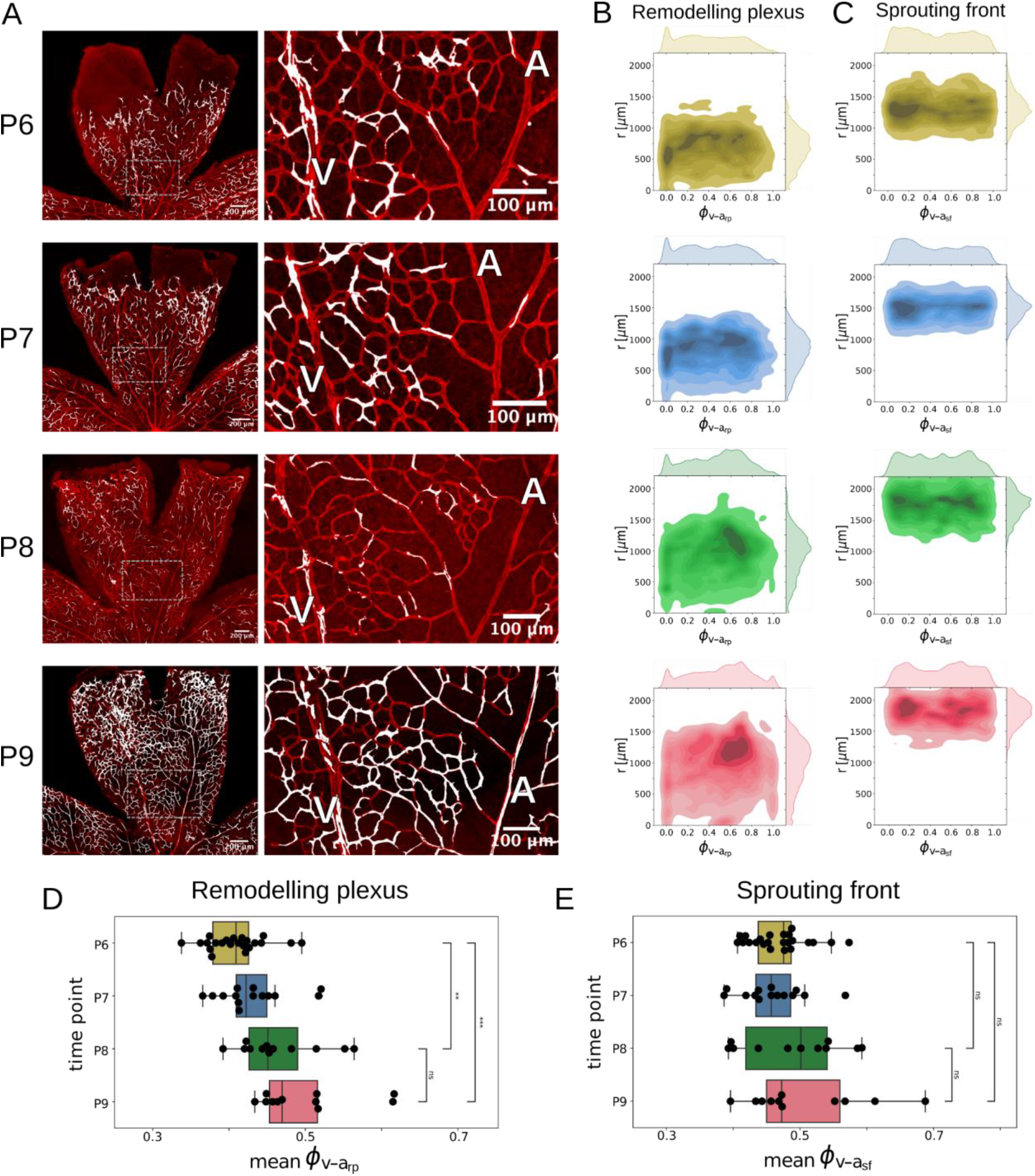
Collective endothelial cell distribution shift from veins to arteries in control retinas. **A**: Examples of retinal images at P6, P7, P8 and P9 and mosaic expression of Cre linked to GFP under a Vegfr3 promoter. While for early time points GFP positive cells are only found in veins and the remodelling plexus, they are also present in arteries for P9 retinas suggesting a movement of ECs from veins to arteries. **B, C:** The endothelial cell distribution is visualized by a kernel density estimate (KDE) plot for each time point in the sprouting front and the remodelling plexus (P6: N=22; P7: N=14, P8: N=12, P9: N=11). **D, E:** For each retina sample (black dots) the mean of the EC distribution in the remodelling plexus along the vein-artery coordinate ϕ_v-a,rp_ is shown. We observed a significant population shift of the labelled EC population towards arteries over time comparing P6 to P8 (p < 0.01) and P6 to P9 (p< 0.001), but not for the last time interval comparing P8 to P9 (p = 0.17). In the sprouting front, we did not observe any significant changes in progression indicated by the mean ϕ_v-a,sf_ of the EC population towards arteries comparing P6 to P8 (p = 0.44), P6 to P9 (p = 0.20) and P8 to P9 (p = 0.66). Statistical analysis was performed using Welch’s t-test.

**Box Figure 1:**
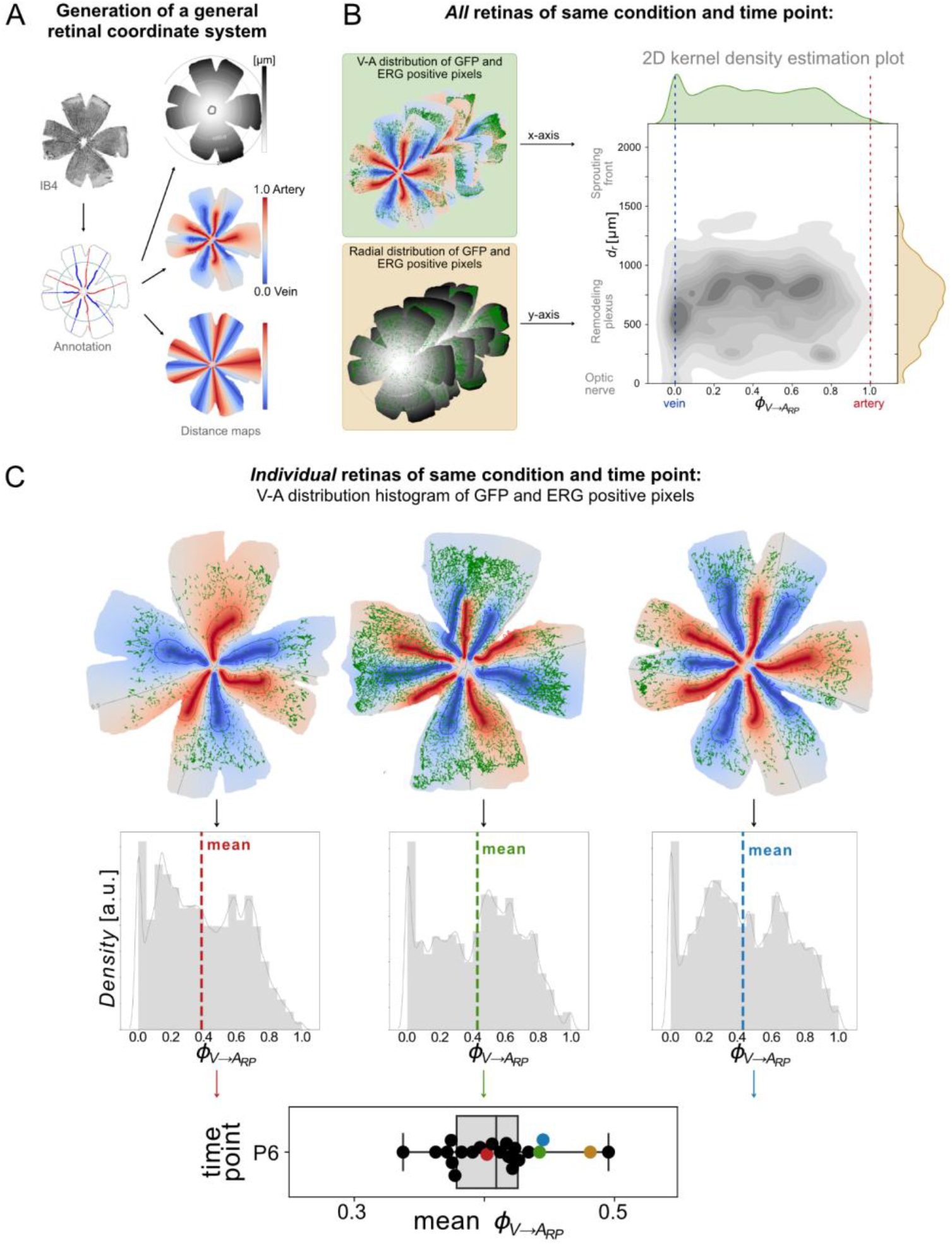
Explanation of the computational analysis method to visualise and quantify the spatial EC distribution in the developing mouse retina. **A** IB4 stainings *(top left)* are used to manually annotate the vascular anatomy of each retina *(bottom left)*: arteries (red), veins (blue), optic nerve (white), ellipse separating sprouting front and remodelling plexus (turquoise), and retina outline (grey). These annotations are the base for distance maps that act as general coordinate systems of each retina: distance to the optic nerve d*_r_ (top right)*, relative distance between veins and arteries ϕ_*v*−*a*,*rp*_*(middle right,* used for analysis in the remodelling plexus), and relative distance between regions in the retina that are more likely to feed into an artery versus regions that are more likely to feed into a vein ϕ_*v*−*a*,*sf*_ *(bottom right,* used for analysis in the sprouting front). The relative vascular distance in the remodelling plexus ϕ_*v*−*a*,*rp*_ was computed using the distance to the nearest artery (*d_a_*) and distance to the nearest vein (*d_v_*) as follows: ϕ_*v*−*a*,*rp*_ = *d*_*v*_/(*d*_*a*_ + *d*_*v*_) such that pixels lying on a vein have a value of 0 and pixels on an artery have a value of 1. The relative vascular distance in the sprouting front ϕ_*v*−*a*,*sf*_ was computed in a similar fashion but using the straight-line vessel extensions for reference. **B** GFP-labelled ECs from all retinas of the same condition are mapped onto the coordinate systems to extract their spatial distribution with respect to the vasculature (ϕ_*v*−*a*,*rp*/*sf*_) and proximity to the optic nerve (*d_r_*). The spatial distributions of several retinas are combined in a 2D kernel density estimation (KDE) plot where darker colours mean higher EC density. The x-axis shows ϕ_*v*−*a*,*rp*/*sf*_ and therefore the relative distance of ECs between veins and arteries. The y-axis shows *d_r_* and therefore the radial position of ECs from proximal (bottom) to distal (top). **C** For statistical analysis, the means of the ϕ_*v*−*a*,*rp*/*sf*_ or *d_r_* of individual retinas of the same condition are plotted in a boxplot.

In order to quantitatively assess the direction and speed of this movement, we developed a coordinate system that quantifies the cumulative distribution of GFP-positive ECs in the retina relative to veins and arteries and the optic nerve. The identity of ECs was verified by using the endothelium-specific transcription factor ERG (ETS related gene) to label endothelial nuclei (Birdsey et al., 2008; Franco et al., 2015). The outline of the retina, optic nerve, arteries and veins are used as a mask for each retina as the basis of a coordinate system (Box Figure 1A). The coordinate system locates any point in the retinal tissue based on the radial distance from the optic nerve, which is denoted by *d_r_*.. The relative vein-artery distance ϕ_v-a,rp_ is computed from

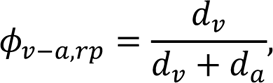

where *d_v_* and *d_a_* are the distances to the closest vein and artery, respectively. The relative distance between veins and arteries ϕ_v-a,rp_ is therefore zero (0) for veins and one (1) for arteries. The mathematically-minded reader will note that this is a curvilinear coordinate system that transforms local and global properties of vascular image data onto a fixed square grid (spanned by *d_r_* and ϕ_v-a,rp_) for computational analysis (Thompson, 1982). Additionally, each retina was divided into a remodelled region containing mature veins and arteries (in the following called remodelling plexus) and a region lacking mature vessels (called sprouting front) by an elliptical domain. To quantify the EC distribution in the sprouting front, we extended the reference system by extending veins and arteries with straight lines into the sprouting front, the corresponding vein-artery distance is denoted by ϕ_v-a,sf_. In this way, we could track EC distribution dynamics separately in the remodelling plexus and the sprouting front. We quantified the cumulative distribution of GFP- and ERG-positive pixels in both reference systems per condition over time and visualized it in 2D kernel density estimation (KDE) plots (Box Figure 1B). For statistical analysis, we plotted the mean of the distributions of individual retinas of the same condition in a boxplot (Box Figure 1C). The distribution of GFP- and ERG-positive pixels serves as a proxy for the distribution of GFP-positive ECs. For the sake of brevity, we will occasionally refer to GFP-positive ECs as “labelled ECs”.

Computation of EC distributions over time revealed a gradual shift of labelled ECs from veins to arteries. Example images of retinas for each time point are shown in Figure 1A. At P6, there are almost no labelled ECs in the artery, whereas at P9 the artery is densely populated with labelled ECs. Computational kernel density estimate (KDE) analysis confirmed the progressive shift of labelled ECs from veins to arteries in the remodelling plexus derived from N=11 to N=22 retina samples (Figure 1B). The shift was significant comparing the mean vein-to-artery distances, ϕ_v-a,rp_, from P6 to P9 and from P6 to P8 (Figure 1D). Conversely, in the sprouting front, we did not observe any significant changes of the mean vein-to-artery distances ϕ|_v-a,sf_ (Figure 1C, E). To obtain a more detailed picture, we also quantified the 10th and 90th percentiles of vein-artery distances ϕ_v-a,rp_ in the remodelling plexus, which describe the accumulation of the labelled EC population near the veins and arteries, respectively (Figure 1 Supplement 1). We observed a significant shift of both the 10th and 90th percentile in the remodelling plexus from P6 to P9. Quantification of the 10th and 90th percentiles of the population distributions also indicate that ECs entered the artery only at later time points, see Figure 1 Supplement 1 A, C.

To understand population movements across the entire retinal segment, including movement between the remodelling plexus and sprouting front, we quantified the radial and vein-to-artery population dynamics in the whole retina by pooling labelled ECs in both the remodelling plexus and in the sprouting front together (Figure 1 Supplement 2). Despite the different EC behaviour in the remodelling plexus and the sprouting front, we found that the overall distribution of labelled ECs progressed significantly shifted towards the arteries from P6 to P9 for the 10th, 50th and 90th percentiles. For the radial movement towards the sprouting front, we observed a significant movement of labelled ECs from P6 to P8 for the 10th, 50th and 90th percentiles.

Our radial analysis revealed that only labelled ECs within the 10th percentile in the radial direction (closest to the optic nerve) show a slight non-significant movement back towards the optic nerve from P8 to P9. This can be explained by labelled ECs entering the artery and moving against the flow, which is directed away from the optic nerve. In the P9 KDE plot in Figure 1B, this phenomenon is visible as a subpopulation of ECs forming a “side arm” from the population bulk, which is migrating along the artery. In summary, our population migration analysis suggests that ECs show a heterogeneous migration pattern, with a tendency for ECs to migrate towards the sprouting front in the whole remodelling plexus except for the artery.

### Computational simulation predicts that EC migration is governed by a dual force-field shaped by VEGFA gradients and shear stress

In order to gain insight into the driving forces underlying these observed population shifts, and to estimate the migration routes and speed, we developed a computational model of stochastically labelled EC agents based on a real segmented retina vein-plexus-artery sector. EC agents were randomly seeded resembling the experimentally observed distribution observed at P6 (Figure 2A), which served as the initial state for all following simulations.

**Figure 2:**
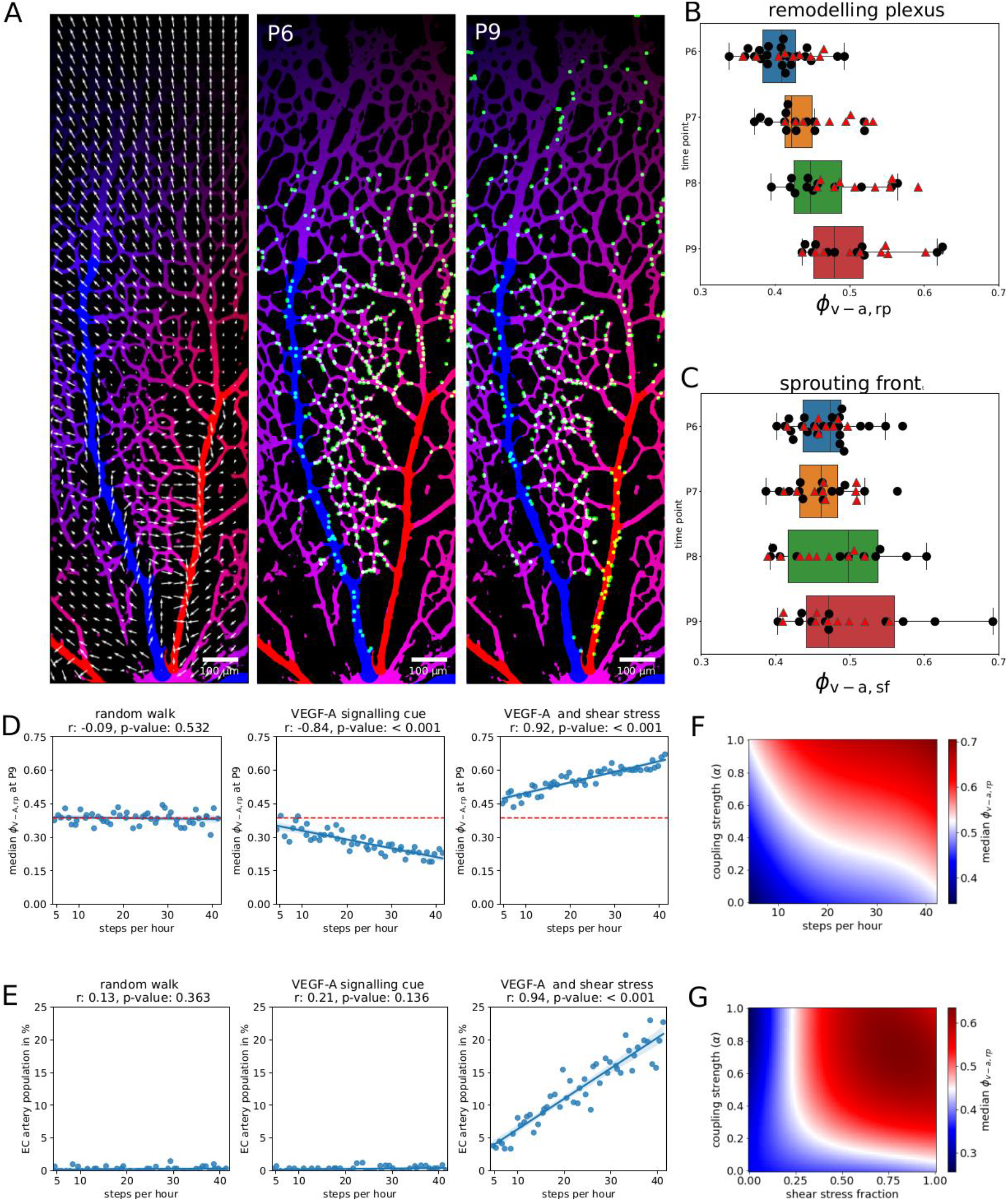
Simulation of EC migration in the retinal vasculature. **A**: Snapshots of the guiding force-field comprising the VEGFA cue and shear forces [*left*] at P6 in a real segmented retina vein-plexus-artery sector derived from image data. Simulated VEGFR3CreERT2;mTmG cells [*right*] at P6 and P9. A substantial fraction of control EC agents have entered the artery at P9. **B:** Comparison of real data (black dots) and model simulations (red triangles). The mean of vein-artery distances in the remodelling plexus (ϕ_v-a,rp_) and **C**: in the sprouting front (ϕ_v-a,sf_), for the best fit μ_*r*_ =18 h^-1^, coupling strength α = 0.36 and fraction of shear stress ξ = 0.53. **D**: Random walk model (M1), VEGFA cue model (M2) and the model incorporating both VEGFA cue and shear stress (M3). Simulated median vein-artery distances ϕ_v-a,rp_ at P9 are plotted against movement rate, red line indicates mean of initial ϕ_v-a,rp_ at P6. No significant correlation of movement rate and ϕ_v-a,rp_ was observed for the random walk model M1 (parameters α = 0.0 and ξ = 0.0), Pearson’s R: −0.09, p = 0.532 (Wald’s test). For model M2 (parameters α = 0.5 and ξ = 0.0), the movement rate and ϕ_v-a,rp_ are negatively correlated with significant population shift towards veins, Pearson’s R: −0.84, p < 0.001 (Wald’s test). Only for model M3 (parameters α = 0.5 and ξ = 0.5), movement rate and ϕ_v-a,rp_ are positively correlated with significant population shift towards arteries Pearson’s R: 0.92, p < 0.001 (Wald’s test). **E:** Dependency of the fraction of EC agents that reached the artery on the movement rate. A substantial population reaches the artery only for model (M3), while almost no EC agents reach the artery for model (M1) or (M2). **F:** Phase diagram for the dependency of vein-artery distance ϕ_v-a,rp_ at P9 on coupling strength (α) and movement rate. **G**: Phase diagram for the dependency of vein-artery distance ϕ_v-a,rp_ at P9 on coupling strength (α) and shear stress fraction (ξ).

We investigated three competing models: The null model (M1) assumes random walk of EC agents in the vascular network. The second model (M2) incorporates the well-known mechanisms of chemoattraction of EC towards VEGFA produced in a graded manner by hypoxic cells in the retinal periphery but does not include mechanical shear stress cues. The third model (M3) assumes, in addition to M2, a physical cue which is a result of blood flow induced shear stress. The shear stress cue is computed from the gradient of the vein-artery distance map, and always points against the direction of blood flow. It is therefore directed from veins to arteries in the remodelling plexus and sprouting front. In labelled arteries, the shear stress field is directed towards the optic nerve and in veins away from the optic nerve. See Supplementary Text S1 for a detailed mathematical description. The force field combining chemo-attractive and physical cues is shown in Figure 2A.

The EC agents are modelled as time-continuous walkers on a lattice and simulated using the well-established Gillespie algorithm (Gillespie, 1977; Giuggioli et al., 2013; Montroll and Weiss, 1965). The waiting time for each agent to move from one lattice site to another is exponentially distributed with mean μr, which is referred to as the movement rate. We introduce a dual force-field 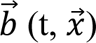, which imposes a bias on the movement of each EC agent and depends on time t and the local position 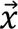 in the vascular bed. This dual force-field is composed of a shear stress dependent component 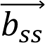 and a VEGFA dependent component 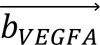 as follows:

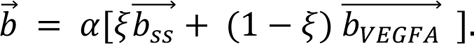

The parameter *ξ* modulates the contribution of the shear stress and the VEGFA component. For *ξ* = 0, the force-field depends entirely on the VEGFA cue, whereas for *ξ* = 1, the force-field depends only on the shear stress component. The parameter α modulates the overall strength of the dual force-field. For α = 0, the EC agents exhibit random walk movement, while for α = 1, the EC agents are strictly coupled to the local force field 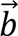 and the movement rate becomes a velocity. In summary, the movement of EC agents in the vascular bed is driven by a dual force-field that can be modulated in strength and composition towards shear stress or VEGFA dependence by the two parameters α and *ξ*.

Next, we sought to understand how random walk, local VEGFA and shear stress components affect the collective population behaviour of ECs in the vascular bed by varying the movement rate in all three models. Does random walk lead to a population shift and if so, in which direction? What is the role of the VEGFA and the shear stress components in driving the EC population towards arteries or veins? Can the increase of the labelled EC population in the artery as observed in the *in vivo* data be explained without a shear stress component? Therefore, we varied the movement rate by an order of magnitude from 4 to 40 steps per hour with a step size of 0.69 μm. This range corresponds to a diffusion constant of 0.5 μm^2^/h to 5 μm^2^/h for random walk movement. For strict coupling to the force field, and therefore directed movement, this range corresponds to a maximum velocity of between 2.76 μm/h and 27.6 μm/h. These boundaries are in the physiological range of measured EC migration speeds *in vivo* (Rosa et al., 2022). The mean vein-artery distance shift in the simulated EC distributions was slightly positive for model M1, but showed no significant correlation with movement rate (Figure 2D, E).

In model M2, the mean vein-artery distance of the EC population correlates negatively with the movement rate in the remodelling plexus, meaning that the EC population shifts towards veins contrary to what was observed in the *in vivo* data. We also found that in the case of models M1 and M2, EC agents almost never reach the artery, even at high movement rates. Only for the model M3, which incorporates both VEGFA and shear stress cues, we observed a significant correlation of movement rate with both observables, the mean population shift in vein-artery distance and the proportion of ECs in the artery. In order to understand the general model behaviour and dependence of the expected vein-artery population shift on key variables, we constructed two phase diagrams by varying the parameters (i) coupling strength and movement rate as well as (ii) coupling strength and proportion of the shear stress. From these phase diagrams we can read off how coordinated parameter changes maintain or change the median vein-artery population shift, see Figure 2F, G.

There are two main aspects that explain this model behaviour. First, the vasculature in the remodelling plexus has an asymmetric hierarchically branched structure with arteries feeding few arteriolar branches, but increasingly more capillary branches that densely feed into the vein. Since the arteriolar branches are less numerous than capillaries, the probability of encountering a random walker is much lower in those arteriolar branches that the EC agents must traverse to reach the artery. Physicists will likely recognise this process as a variant of the narrow escape problem (Grebenkov et al., 2019; Holcman and Schuss, 2014), where the escape time of a random walker (here escape of ECs from the remodelling plexus into the artery) is highly dependent on the geometry of the domain (plexus) and size of the exit sites (here arteriolar branches of the artery). In the case of the VEGFA cue pointing towards the sprouting front, this encounter frequency in arteries is further reduced by the orientation of the movement away from the artery and its side branches.

Second, random walk is an inefficient means of transport compared to directional movement. This can be explained by the expected distance travelled: the root-mean square displacement scales with 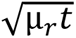 in the case of random walk (α = 0), while in the case of strong coupling (α = 1) it scales with μ_*r*_*t*. This means a 10-fold increase in the movement rate results in an increase by a factor of about 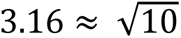 of the expected net distance travelled, while for strict coupling and directed movement the net distance travelled increases by a factor of 10 (Codling et al., 2008).

The phase diagram further suggests that directional movement is highly robust over a range of movement rates as long as the coupling strength is above α = 0.2 (Figure 2 F). In biological terms, this means that even a rather weak 60/40 bias in directionality towards arteries is sufficient to drive the observed population shift. Furthermore, the phase diagram predicts that at higher coupling strengths, a slow movement rate is sufficient. This could mean that at high shear stress values, where the directionality stimulus is very pronounced, slow movement would be sufficient, whilst a weak directionality stimulus requires higher movement rates to be effective. It should be noted that a high movement rate at low coupling of the EC population can be interpreted as increased heterogeneity in the EC population meaning that not all cells are following the same cues.

To estimate EC migration velocities in the vascular bed, we first needed to find optimal model parameters for the relative coupling strength to shear stress and VEGFA signalling based on the experimental data. To do so, we performed a parameter scan of the coupling strength (α) to the force field and the proportion of coupling to the shear stress cue (ξ) using Gaussian process regression (Mirams et al., 2016) to effectively obtain parameters that matched the experimentally observed data (see parameter values in Table S1). Conclusions are based on the degree of similarity between the measured and simulated mean, median, 10th and 90th percentile of vein-artery and radial distances of the EC distribution in the remodelling plexus and sprouting front. Since, as mentioned above, a weak coupling to the force-field can be partially compensated by a higher movement rate, we set the movement rate at μ_*r*_ =18 steps per hour. This movement rate corresponds to a maximum migration speed of 12.42 μm/h, assuming that the EC agent moves constantly in one direction, which is in the range of experimental observations *in vivo* and *in vitro* (Rosa et al., 2022; Weijts et al., 2018). Parameter inference yields an overall average coupling strength of α = 0.36 with a fraction of the shear stress component of ξ = 0.53 (see Table S2). The proportion of the VEGFA component is given by 1 −ξ = 0.47, which is slightly lower than the shear stress component. These results of the parameter scan predict that the observed movement of ECs *in vivo* can be explained by a dual force-field in which VEGFA and shear stress components contribute approximately equally to the dual force field, but with a slight bias towards the influence of shear stress. A comparison of simulated vein-artery distances in the remodelling plexus and sprouting front is shown in Figure 2 B, C.

We further asked how these model parameters translate into actual migration speeds in the vascular bed. Example trajectories in the remodelling plexus and sprouting front are shown in Figure 2 Supplement 1 A. We ran a series of N=10 simulations to measure the resulting migration speeds. In both the remodelling plexus and the sprouting front, the average speed is approximately the same with a mean of 2.50 μm/h and 2.53 μm/h, respectively. However, their directionality is entirely different. Labelled ECs in the remodelling plexus are migrating towards the artery at a speed of 0.53 μm/h, while labelled ECs in the sprouting front are moving away from the reference artery with an average speed of 0.59 μm/h. Note that without any bias towards artery or vein, migration in the sprouting front away from the optic nerve results in larger distances towards the reference arteries and veins. Therefore, a net migration away from the artery is expected in the sprouting front. This behaviour reveals an interesting migration route. A part of the EC population migrates towards arteries but eventually enters the sprouting front to compensate for a population shift away from arteries induced by the VEGFA gradient. The other part of the EC population enters the artery and migrates towards the optic nerve. This is illustrated by the bidirectional migration pattern of ECs visible in the trajectory plot Figure 2 Supplement 1A.

Finally, an examination of the relationship between the relative weight of VEGFA and shear stress influence (termed the shear stress fraction) and the overall coupling strength α, revealed an exquisite sensitivity of ECs to the shear stress cue (see phase diagram, Figure 2G). At very low overall coupling to the force field (α below 0.1), the relative effect of shear stress would need to dominate in order to explain the observed EC population shifts towards arteries (Figure 1). However, already at coupling strength of α = 0.2 (meaning that EC movement is heterogeneous and has a major random component), even a minor influence of shear stress compared to VEGFA cue, is sufficient to explain the population shift towards arteries.

### Cdc42 is essential for flow-migration coupling and EC migration to arteries

Cell migration processes are primarily driven by actomyosin and microtubule dynamics, which are regulated by actin nucleators under the control of small Rho GTPases (Tzima et al., 2003; Zegers and Friedl, 2014). The ability of cells to move individually or collectively relies on dynamic protrusion cycles of leading membrane structures, such as lamellipodia and filopodia, as well as the formation of new adhesion structures along these protrusions to exert the force that drives forward movement. Unlike filopodia, which extend in multiple directions in migrating cells and are formed by linear actin fibres, lamellipodia are formed by branched actin networks. Stress fibres that run along the cell axis are connected to integrin-based adhesions. A large body of work has established that the small Rho GTPase family members Cdc42 and Rac1 differentially drive the formation of lamellipodia and filopodia, respectively (Etienne-Manneville and Hall, 2002; Lawson and Ridley, 2018; Raftopoulou and Hall, 2004; Ridley, 2001). Depletion of either of these in ECs *in vitro* has been reported to interfere with EC migration. *In vivo*, deletion of either Cdc42 or Rac1in ECs leads to embryonic lethality (Barry et al., 2015; Tan et al., 2008), and causes retinal vascular patterning defects when deleted during post-natal retina development (Heynen et al., 2013). As vascular patterning defects immediately precipitate hemodynamic changes, the precise role of a given Rho GTPase in cell-autonomous EC migration behaviour is difficult to disentangle from the confounding influence of patterning changes.

Given the prediction of a dual force-field driving the directional migration behaviour in vascular remodelling, we set out to test the role of Cdc42 in chemotaxis and flow-migration coupling *in vivo*. We used the same labelling technique as in the previous sections with Vegfr3-CreERT2 and R26RmTmG alleles, but in combination with homozygous floxed alleles for Cdc42. We will refer to the EC-specific inducible Cdc42 knockout as Cdc42 iECKO (Figure 3A). Note that the overall retinal vascular patterning remained unaffected, due to the overabundance of unrecombined ECs that remained Cdc42-competent.

**Figure 3:**
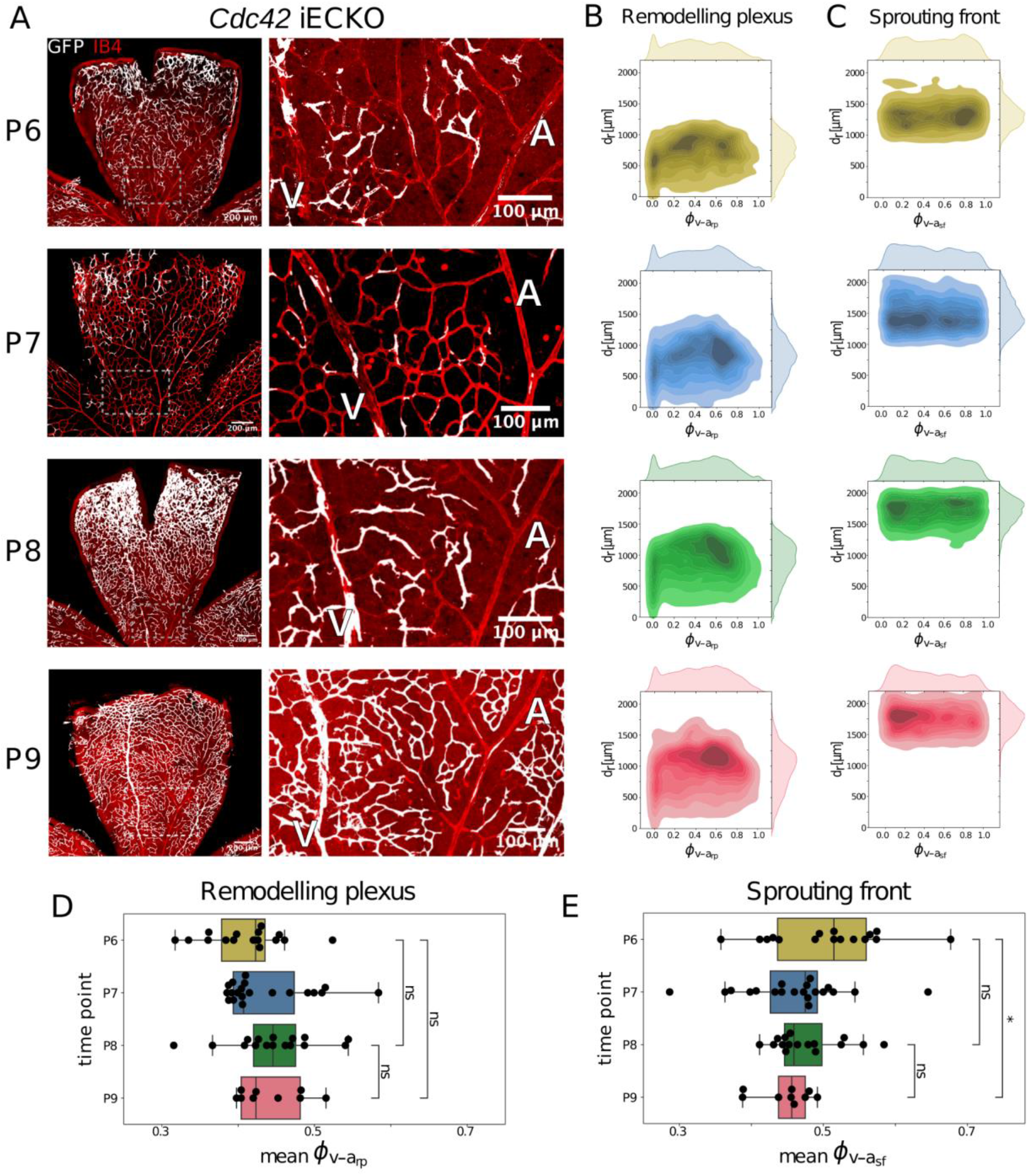
Collective distribution of endothelial cells deficient in Cdc42 stagnates in the remodelling plexus and shifts towards veins in the sprouting front. **A**: Examples of retinal images at P6, P7, P8 and P9 Cdc42 homozygous floxed mice. **B, C:** The endothelial cell distribution is visualized by a kernel density estimate (KDE) plot for each time point in the sprouting front and the remodelling plexus. (P6: N=16; P7: N=20; P8: N=16, P9: N=9). **D, E:** For each retina sample (black dots), we computed the mean of the EC distribution for the remodelling plexus and sprouting front along the ϕ_v-a,rp_ and ϕ_v-a,sf_ coordinate, respectively. In the remodelling plexus, we observed no significant shift of the labelled EC population comparing P6 to P8 (P = 0.09), P6 to P9 (P = 0.16) and P8 to P9 (P = 0.76). However, in the sprouting front we observed a significant shift of the mean of the EC distribution towards veins comparing P6 to P9 (P < 0.02), but not for P6 to P8 (P = 0.22) or P8 to P9 (P = 0.1). Statistical analysis was performed using Welch’s t-test.

Quantification of labelled EC populations in our vein-to-artery coordinate system over the same developmental stages revealed a profound difference in the evolution of EC distributions over time upon Cdc42 depletion (Figure 3B,C) compared to control (Figure 1 B,C). Detailed comparison of vein-to-artery distances of P6, P7, P8 and P9 retinas illustrated a minor shift of labelled EC populations in the plexus towards the centre between artery and vein until P8, but no further shift thereafter until P9 (compare Figure 3B, D). Remarkably, even in retinas with a high recombination frequency (example shown for P8 and P9 in Figure 3A), the artery remained almost completely free of labelled ECs. These results demonstrate that ECs deficient in Cdc42 fail to effectively move towards and into arteries. Furthermore, Cdc42 depleted ECs get stuck in the vein with no significant change of the 10th percentile in the vein-to-artery distance over the whole time period (Figure 3 Supplement 1A).

Statistical comparison of the distributions showed that cells in the sprouting front shifted towards the region peripheral to the forming vein (Figure 3E). This could be driven by local proliferation, known to be highest in that domain. Note that ECs depleted of Cdc42 proliferate at normal rates (Laviña et al., 2018). Alternatively, the net movement of non-recombined ECs could cause an apparent “backward” shift, even if the Cdc42 depleted cells are not actively migrating.

Surprisingly, however, Cdc42 depleted ECs continued to migrate towards the retinal periphery in the sprouting front (Figure 3C, E), providing an intriguing third possibility: Cdc42 is exquisitely necessary for cells to move against the shear gradient under flow, but less so for chemotaxis driven by VEGFA gradients. In fact, this could be confirmed by parameter inference of the computational model, when applied to the experimental data from Cdc42 mutant cells. Whereas the overall strength of the movement bias, indicated by the coupling strength α, is slightly reduced in Cdc42-depleted cells from α =0.36 (control) to α = 0.26, the proportion of the shear stress cue is reduced by more than two-fold from ξ =0.53 (control) to ξ = 0.19 in Cdc42-depleted cells. However, the coupling to the VEGFA cue appears to be almost unaffected with α_VEGFA_ = α (1-ξ) = 0.21 compared to α_VEGFA_ = 0.17 in control.

We further tested how these model parameters changed the actual migration velocity estimates. Despite the differences in coupling strength, the total velocity of Cdc42 depleted ECs was similar to the control in the sprouting front and slightly higher in the remodelling plexus (Figure 3 Supplement 3B). This counterintuitive prediction can be explained by two effects: First, the shear stress and VEGFA cues are additive in the vein, meaning that both cues point in the same direction, while closer to the artery these cues are competing, and (2) the vein is a comparatively straight vessel, while migrating through the maze-like capillary network results in longer distances and therefore shorter net displacement. Both effects cause ECs that remain in or close to the vein to have higher net displacement.

Indeed, in the remodelling plexus, the net migration of Cdc42-depleted cells towards the artery is zero (Figure 3 Supplement 3C). Even though there is a slight coupling to the shear stress field, the net migration is counteracted by the VEGFA cue and a substantial fraction of Cdc42 depleted cells follow the route along the vein as discussed above. In the sprouting front, Cdc42-depleted ECs migrate away from the reference artery, as they follow the route to the sprouting front along the VEGFA gradient.

### Rac1 deficiency causes minor defects in directional migration

The small Rho GTPase Rac1 has long been recognized as essential driver for invasive cell behaviour, including in collective cell migration scenarios, both in developmental tissue morphogenesis (Tan et al., 2008), as well as in cancer (Liang et al., 2021). It is best known for a key role in promoting branched actin structures during lamellipodia formation. Rac1 activation is essential to provide spatial information for shear stress-induced cell alignment and is furthermore regulated by shear-induced new integrin binding to the extracellular matrix (Tzima, 2002). We therefore tested the particular role of Rac1 in endothelial flow-migration coupling and chemotaxis using the same approach as for Cdc42 (Figure 4A). We will refer to the EC-specific inducible Rac1 knockout as Rac1 iECKO.

**Figure 4:**
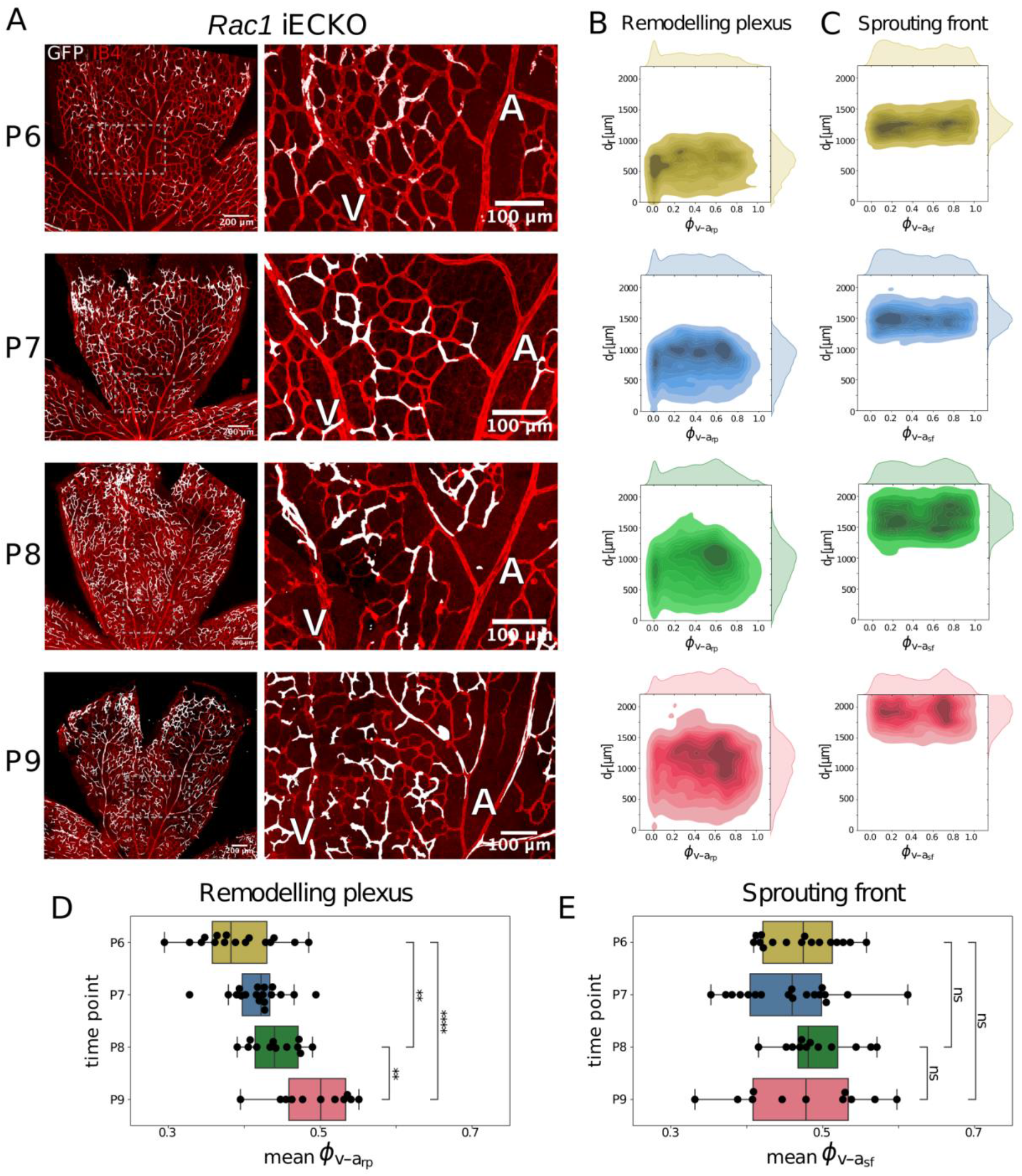
Collective population shift of endothelial cells deficient in Rac1 reveals mild migration defects. **A**: Examples of retinal images at P6, P7, P8 and P9 Rac1 homozygous floxed mice. **B, C**: The endothelial cell distribution is visualized by a kernel density estimate (KDE) plot for each time point in the sprouting front and the remodelling plexus (P6: N=16; P7: N=18; P8: N=12, P9: N=11). **D, E**: For each retina sample (black dots), the mean of EC distribution along the ϕ_v-a,rp_ and ϕ_v-a,sf_ coordinate is shown in the remodelling plexus and sprouting front, respectively. In the remodelling plexus, we observed a significant shift of the labelled EC population towards arteries comparing P6 to P8 (p < 0.01), P6 to P9 (p < 0.001) and P8 to P9 (p < 0.01). However, in the sprouting front, we did not observe any significant shift of the mean of the EC distribution comparing P6 to P8 (p = 0.17), P6 to P9 (p = 0.83) and P8 to P9 (p = 0.53). Statistical analysis was performed using Welch’s t-test.

Unlike Cdc42 depleted cells, Rac1 depleted cells still showed significant displacement towards the artery in the remodelling plexus, albeit with slower initial progression (Figure 4 B, D) compared to control (Figure 1 B, D). Quantitative comparison of the mean vein-to-artery distances in the remodelling plexus even revealed a conspicuous acceleration of the population shift between P8 and P9. Nevertheless, only few cells reached the arteries by P9.

The sprouting front showed no significant defect in the migration of Rac1 depleted ECs, further suggesting that Rac1 is less important for VEGFA chemotaxis. In particular, we observed that EC migration towards the sprouting front is not delayed from P6 to P8 in contrast to the vein-to-artery migration (Figure 4 Supplement 2 D, E, F). Consequently, there is an increased shift of the labelled EC population towards the sprouting front compared to control. Interestingly, this may also explain the observation that we did not find a significant vein-to-artery shift of the total labelled EC population in the whole retina (Figure 4 Supplement 2 B).

Estimation of the computational model parameters to replicate the experimental *in vivo* data confirmed these observations. The directionality of movement in the case of Rac1 depleted cells is almost the same (α = 0.32) compared to control (α = 0.36), while the coupling to the shear stress cue is reduced from ξ =0.53 (control) to ξ = 0.34. This indicates a shift in the modulation of the dual force-field with less sensitivity to shear stress and more sensitivity to the VEGFA signalling cue than in control.

We further tested how these model parameters for Rac1 depleted ECs translate to actual velocity estimates. In the remodelling plexus, total velocities are slightly higher for Rac1 depleted ECs compared to control, but not in the sprouting front (Figure 4 Supplement 3B). At the same time, net migration towards arteries (Figure 4 Supplement 3C) is slightly slower than control. This effect is again explained by the geometry of the vascular network (straight vein versus maze-like network) and the fact that shear stress and VEGFA cues are additive in the vein, while they are competing closer to the artery and in the remodelling plexus. The speed-up of migration that we observe *in vivo* is not well captured by the model, since the coupling strength is not changing over time as well as the coupling to the shear stress field, but this may be the case during vascular remodelling. Despite this limitation, the data and model suggest a slightly perturbed migration route in which Rac1-depleted ECs are initially more likely to migrate towards the sprouting front and only migrating from the remodelling plexus towards the arteries at later time points in development.

### Cdc42, but not Rac1, drives polarized flow-migration *in vitro*

We found impaired vein-to-artery migration in Cdc42-depleted ECs and also accelerated vein-to-artery migration of Rac1-depleted cells from P8 to P9 compared to control *in vivo* (see previous sections). We therefore investigated the role of Cdc42 and Rac1 in EC migration behaviour under controlled *in vitro* conditions. To assess EC migration under coupling to shear forces *in vitro*, siScrambled (siScr) control, siCdc42 and siRac1 treated endothelial monolayers were exposed to 20 dynes/cm2 of flow using the Ibidi perfusion system and observed for up to 17 hours using a non-toxic fluorescent DNA dye for nuclear tracking. Following acquisition of time-lapse image sequences, the trajectories of all individual cells were obtained using TrackMate (Ershov et al., 2022). Individual cell trajectories from example movies from each condition are shown in Figure 5A after 2h of flow exposure. For all conditions, there is an overall tendency of cells to migrate against flow. Strikingly, this tendency was notably reduced in the siCdc42 condition compared to siScr and siRac1, suggesting a role for Cdc42 in aiding ECs to migrate against flow. In contrast, in the siRac1 condition, cell migration was on average more directed against flow than in siCdc42 and even slightly more directed than in siScr. However, siRac1 treated cells migrated at an overall slower speed than siScr, which together results in a similar net displacement of ECs with respect to the flow direction (Figure 5A).

**Figure 5:**
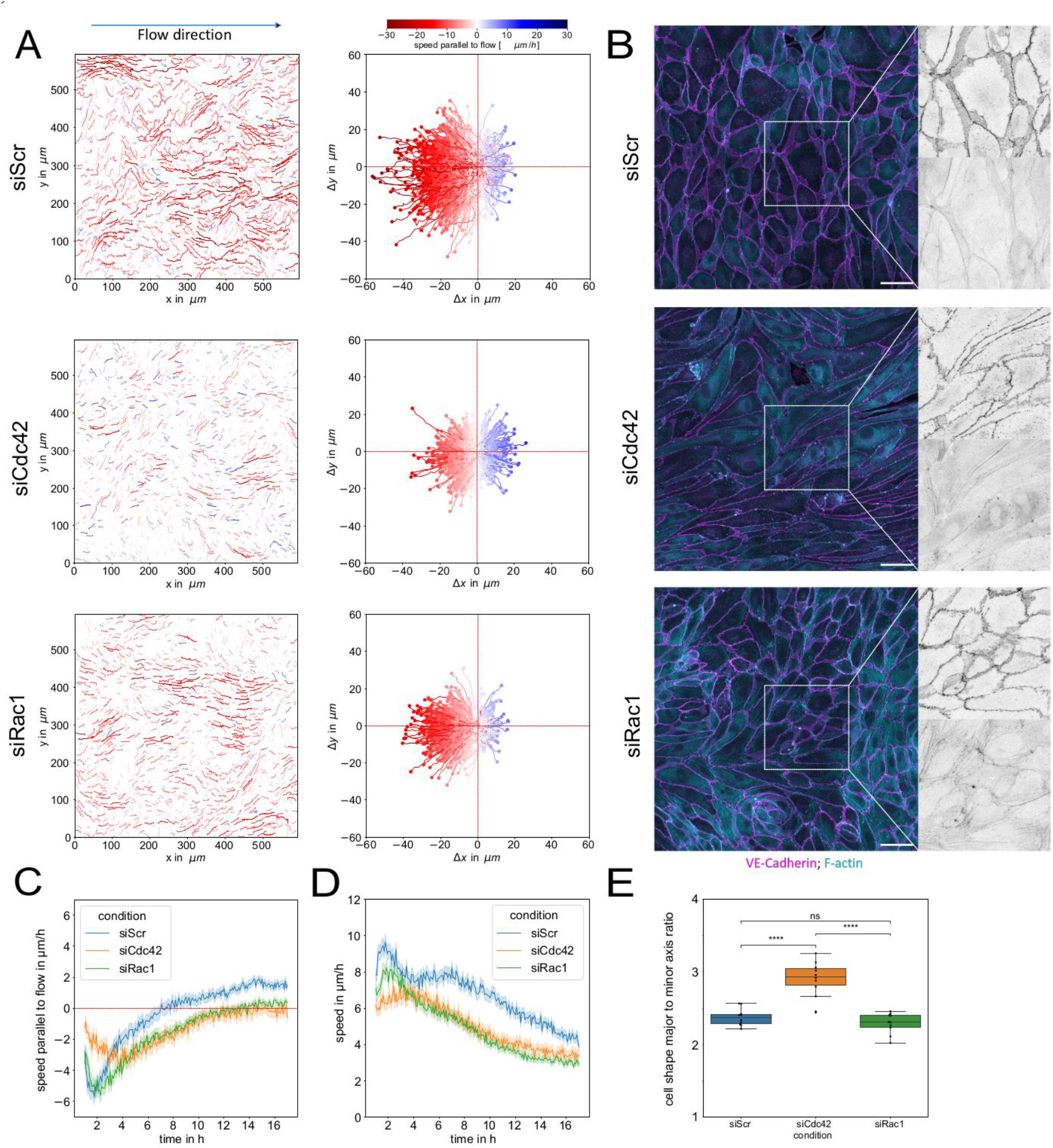
Migration impairment following Cdc42 and Rac1 loss-of-function in endothelial cells. Directional migration *in vitro*: **A:** Cell trajectories computed from fluorescence images of HUVEC nuclei following siScrambled (siScr), siCdc42, and siRac treatments, during exposure to arterial high shear stress (20dyne/cm^2^) at the 2h time-point. On the left side, the cell trajectories are shown with their original spatial position in the time-lapse movies, while on the right side the same trajectories are plotted with the initial coordinates of each cell being set to 0. **B:** Immunofluorescence images of HUVEC monolayers following siScr, siCdc42, and siRac1 treatments. Cells were fixed after the migration assay at t=17 h and stained for VE-Cadherin (magenta) and F-Actin (cyan), scale bar: 50 µm. White dashed rectangles show area of zoomed in individual channels on the right. **C:** Time series depict the progression of cell speeds parallel to flow under the different knockdown treatments over a period of 17 hours. Negative cell speeds correspond to migration against flow. In siScr, a higher cell ratio moves against the flow. This tendency is reduced in siCdc42, while under siRac1 conditions, a larger proportion of cells move against the flow, with trajectories showing a more horizontal orientation. **D:** Time series of total velocities, which are computed from the total net displacement, irrespective of migration direction. **E:** Major/minor axis ratio cell shape quantification of HUVECs after 17h of exposure to flow, n=10 images per condition.

Further quantification of the cell speeds over the entire time series (Figure 5C) corroborated these observations. ECs treated with siCdc42 exhibited a significantly slower peak velocity against flow compared to both siScr and siRac1 conditions. The total migration speed without considering the direction is significantly larger for siScr than for the other conditions (Figure 5D). Following a strong initial migration response of siScr treated cells against flow, after 8 hours ECs displayed an adaptation phase with slow or zero net migration in flow direction (Figure 5C). This adaptation was considerably delayed for siRac1 and siCdc42 (Figure 5C). At the same time, total speed of migration decreased much stronger for siCdc42 and siRac1 than for siScr over the whole time period (Figure 5D). In summary, these results suggest an impairment of the migration apparatus upon treatment with both siRac1 and siCdc42, with Cdc42 being particularly responsible for the impairment in the initial flow-migration coupling.

Immunostaining of each condition revealed striking differences in cell shape, junction formation and cytoskeletal organisation. In the siCdc42 condition, cells appeared strongly elongated parallel to the direction of flow (Figure 5B). Interestingly, siCdc42 cells showed defective junction formation (VE-cadherin staining), with long junctions parallel to flow that seemed unable to stabilise into linear cell-cell connections, and a reduced presence of actin cables at junction sites. These observed characteristics are consistent with previous theoretical studies, where such cellular features are indicative of a solid-like behaviour or jammed state, implying that siCdc42 ECs would move slowly only in one direction while lacking the ability to rearrange (Bi et al., 2015; Lenne and Trivedi, 2022; Trepat and Sahai, 2018).

To understand the role of Cdc42 and Rac1 in EC migration independent of flow, we performed live imaging of wound healing assays, as shown in Figure 5 Supplement 1. EC movement was tracked using Hoechst-33342 nuclear labelling. All conditions were imaged simultaneously, starting within 30 minutes of wound-gap formation. Analysis of wound healing kinetics revealed a pronounced delay in wound closure for both siCdc42 and siRac1 conditions. Furthermore, when analysing the migration trajectories of the leading-front ECs within the monolayer (within <200 microns from the wound edge), siCdc42 cells exhibited a significantly reduced total velocity, total displacement (measuring the straight-line distance between the initial and final position) and total distance travelled (quantifying the cumulative path length travelled from the initial to the final position). Similar to the junctional features observed under flow conditions, immunostaining of migrating monolayers consistently revealed discontinuous and weakened cell-cell contacts.

Taken together, these findings highlight the critical role of Cdc42 in facilitating dynamic migration of intercalating ECs, which is required both to effectively close a wound in a monolayer, but more importantly to enable balanced EC arrangements that govern vascular remodelling as the key adaptation mechanism to blood flow. These findings further highlight and confirm that the endothelial junction is the critical mediator of EC rearrangements and that Cdc42 and its effectors are crucial regulators of junctional actin dynamics.

## Discussion

Our innovative combination of stochastic single cell labelling and a bespoke coordinate system to track EC populations across multiple developmental remodelling stages in the retinal plexus identified that ECs originating from the vein and capillary plexus migrate directionally towards the forming arteries. These findings generally agree with previous reports of tip cell to artery movements, and vein to artery migration in development (Barbacena et al., 2022; Lee et al., 2021; Pitulescu et al., 2017; Xu et al., 2014). Until recently, however, the underlying mechanisms, and a quantitative understanding of relative migration speed or detailed routes remained unclear. Barbacena et al. elegantly demonstrated that ECs become polarized by competing forces exerted by blood flow and the chemotactic pull of the VEGFA gradient at the sprouting front. Whereas Barbacena et al. used nuclei-Golgi polarity as a surrogate marker for directional migration, our tracking approach provides detailed population data of migration routes. Interestingly, although our results confirm the validity of the idea of competing force fields exerted by VEGFA gradients and shear stress, we find that ECs show heterogeneous movements throughout the remodelling plexus, suggesting that this competition (in contrast to the model by Barbacena et al.) does not result from a narrow region of influence of chemokine signalling. Our analysis shows that the transition between VEGFA and shear stress influence is much more gradual regarding migration and changes along the vein-to-artery coordinate. Accordingly, net movement of cells can be observed away from the optic nerve also in the remodelling plexus (though slower), even close to the optic nerve. In contrast, (Barbacena et al., 2022) show that nuclei-Golgi polarity is directed away from the sprouting front and towards the optic nerve in large parts of the retina.

Conceptually, our finding of such heterogeneity is particularly relevant when considering the vein, and its proximal plexus. If all ECs were to strictly follow the VEGFA and shear stress gradients in the vein, where these force fields notably align, unlike in the remodelling plexus or in the artery, ECs would rarely be able to leave the vein. Instead, they would migrate along the vein and towards the sprouting front. Consequently, this would lead to proximal rarefaction of the vasculature in the vicinity of the vein. Thus, our data and model suggest that such heterogeneity in the movement of ECs is important for maintaining a balanced plexus. Computationally, this is modelled as a balance between isotropic movement (or unbiased random movement) and biased movement along the two force fields. Importantly, this convergence of VEGFA and flow-induced migration direction in the vein has not been considered in the analysis by Barbacena et al., which may also explain why the presence and requirement of this heterogeneity was not discovered.

One limitation of our model is that we did not include EC proliferation. Since EC proliferation mainly occurs in the venous bed (Laviña et al., 2018), the overall EC population density would consequently shift towards veins if cells were not leaving and migrating towards arteries. This means that such localised proliferation must be balanced by active migration towards arteries, otherwise the vein would grow dramatically in size. This is precisely what happens in retinas, where most of ECs lack Cdc42, precipitating capillary-venous malformations (Laviña et al., 2018). It also means that our estimates of vein-to-artery migration derived from our model are probably slightly too low, as including proliferation would necessitate faster migration towards the artery to match the observed population shift. A future avenue to consider are the feedbacks of EC migration on the vascular network as described in (Crawshaw et al., 2023; Edgar et al., 2022; Tabibian et al., 2020) and signalling interactions between ECs such as the VEGFA-Delta-Notch pathway (Bentley et al., 2014; Stepanova et al., 2021). In our experimental setup, the fraction of knockout cells was too low to induce significant changes and feedbacks on the vascular network, but with higher knockout frequency this may become an important aspect to consider.

Given the general appreciation of the importance of Cdc42 for cell migration in various cell types and tissues, one of the most remarkable results of our present study is the finding of a very distinct sensitivity of endothelial flow-migration responses to Cdc42 deletion. Migration towards VEGFA and thus towards the sprouting front appeared to be much less affected, despite the prominent role of Cdc42 in endothelial filopodia formation. This intriguing finding was only possible thanks to the combination of dynamic EC distribution analysis and computational simulation, that allowed us to perform parameter scans to extract the relevant coupling strength parameters explaining the observational data.

Why then would migration towards the retinal sprouting front, and thus towards VEGFA be less sensitive to the loss of Cdc42? We previously reported that tip cell guidance towards VEGFA in zebrafish intersegmental vessels continued even in the absence of endothelial filopodia, demonstrating that directional EC migration does not rely on filopodia, one of the key actin-based protrusions that are driven by Cdc42 activity (Phng et al., 2013). This was also recently confirmed by Franco et al. studying the role of Arp2/3 in retinal angiogenesis (Figueiredo et al., 2021). Similarly, a key member of the formin family of actin nucleators was dispensable for tip cell migration towards VEGFA, but essential for endothelial lumen formation and intercalation movements driven by junctional actin polymerisation to stabilise nascent vessels in the fish (Phng et al., 2015). In addition, Cdc42 has been shown to play a critical role in strengthening coupling between adherens junctions and the actin cytoskeleton (Carvalho et al., 2019). The ability of junctional VE-cadherin, the central adhesion molecule of endothelial cell-cell junctions, to bind actin through the catenin complex has been identified as critical for EC intercalation movements in fish (Sauteur et al., 2014).

Accordingly, fish carrying a deletion of the cytoplasmic tail of actin-binding VE-cadherin, showed no defects in sprouting, but failed to stabilise lumenised intersegmental vessels. The authors showed that this defect was due to an inability of ECs to move between each other, as they could not extend their junctional interface in order to intercalate.

We recently identified another actin nucleator required for this endothelial behaviour in zebrafish, the Wiskott-Aldrich syndrome protein Wasp, in particular the fish homologue gene wasb (Rosa et al., 2022). ECs lacking wasb also showed little difficulty in sprouting, but failed to perform intercalation movements that balance out the diameter of veins and arteries. Our time-lapse movie analysis in zebrafish identified that wasb is essential for differential EC movements in future arteries and veins in the trunk of the fish, with ECs failing to move from veins to arteries. This is relevant because the Wasp protein is normally autoinhibited until it is activated by Cdc42. Only activated Wasp can nucleate new actin fibers. Our studies revealed that ECs lacking Wasp failed to nucleate junctional actin fibers (Rosa et al., 2022). Intriguingly, we here find the very same sensitivity of ECs to the function of Cdc42, which is required for EC migration towards the artery *in vivo*, against flow *in vitro*, and also for junctional actin fibre formation. Taken together, these results strongly suggest that endothelial flow-migration coupling, which induces EC migration from the vein to the artery requires a regulated actin-nucleation machinery that drives junctional organisation and dynamics.

VE-cadherin based structures important for cellular rearrangements have been described as junction-associated lamellipodia (JAIL) in endothelial monolayers (Cao et al., 2017) or as junction-based lamellipodia (JBL) in zebrafish *in vivo* (Paatero et al., 2018). Schnittler and colleagues propose that larger JAILs at the polar ends of ECs drive forward movement, whereas smaller lateral JAILs are involved in cellular rearrangements (Cao and Schnittler, 2019). These junctional lamellipodia are formed by Arp2/3 dependent branched actin networks. In line with this notion, Cdc42 and Wasp act upstream of Arp2/3, and both siCdc42 ECs (Figure 5 B) and siWasp ECs conspicuously lack such larger VE-cadherin patches indicative of JAILs (Rosa et al., 2022). Interestingly however, Rac1 has been proposed to be important for JAIL formation (Cao and Schnittler, 2019), yet we find that Rac1 deficiency is less disruptive to flow-migration coupling compared to Cdc42 deficiency. Also, the lack of larger VE-cadherin patches, and the defects in junctional actin organization appear less pronounced in siRac1 ECs, together suggesting that both *in vitro* and *in vivo*, Cdc42 activity is most relevant for junctional organisation and intercalated cell movements within the monolayer or plexus.

While Rac1-depleted cells show only a slight change in migratory behaviour in the initial flow response, the difference is striking at later time points *in vivo*. We observed that the vein-artery population shift was accelerated from P8 to P9 compared to control, suggesting that Rac1 deficiency is potentially more detrimental in low shear conditions, but less so in the high shear regions closest to the artery. Similarly, we observed in our *in vitro* experiments that siRac1-treated ECs continue to migrate against flow, whereas for siScr-treated ECs the net migration against flow equilibrates to zero after approximately 8 hours of migration against flow. This leads us to hypothesise that the Rho GTPases Cdc42 and Rac1 play complementary roles in flow-migration coupling, with Cdc42 being essential for flow sensing and Rac1 being necessary for adaptation to blood flow induced shear stress.

Our trajectory analysis *in vivo* and *in vitro* revealed a surprising heterogeneity in EC movements (Figure 2 Supplement 1 A and Figure 5A), with many moving directionally towards the artery, but also a significant proportion moving towards the sprouting front. This raises the intriguing possibility of regulated cell sorting, where some cells with distinct properties are favoured to reach the artery. Evidence for this idea stems from recent work on coronary vessel formation, which identified so-called pre-arterial cells in the immature plexus, that preferentially form arteries (Su et al., 2018). In addition, the previous finding that arteries are formed by tip cells (Ara et al., 2005; Ivins et al., 2015) proposed that distinct properties including Notch signalling activity, bias ECs to move to the artery. Thus, arterial identity may not be acquired only after ECs have formed a structurally recognisable arterial vessel, but rather arise in cells endowed with specific signalling properties, including Notch, Cx37 (Fang et al., 2017), Cxcr4 (Su et al., 2018), EphrinB2 (Kim et al., 2008), Erk (Deng et al., 2013), that predispose these cells to migrate into a part of the vasculature that will form the future artery. Such a mechanism would also explain how cell sorting via repulsive signalling involving EphrinB2 and EphB4 would work to shape arteries and veins, respectively. Several studies further show that exit from the cell cycle is a necessary driver of arterial cell identity, linked to Notch activity and movement towards the artery (Chavkin et al., 2022; Fang et al., 2017; Luo et al., 2021). Our identification of heterogeneous movement patterns would fit well with the idea of cell sorting, but further suggests that actin regulation through Cdc42 could be a critical effector of pathways governing arterial identity. Conceptually, this would mean that movement heterogeneity involves different levels of Cdc42 activation, with those cells that have more Cdc42 activity making their way towards the artery. Intriguingly, a recent study has uncovered that ECs lacking the SRC family kinase YES migrate preferentially and faster to the artery (Jin et al., 2022), and these ECs actually show higher levels of Cdc42 activity (see Jin et al. supplementary material). Future work will need to specifically test whether such heterogeneity in Cdc42 activity exists, and whether it overlaps with any or all of the above markers or properties of pre-arterial identity.

Addressing such questions has however been extremely challenging in the past, as spatial mapping of single cell data has only recently become sufficiently advanced, but it has been impossible to track populations in dynamic systems. Our development of a bespoke coordinate system and method for tracking populations of stochastically labelled cells, opens new possibilities, as tracking is no longer dependent on unique cell identities. Our coordinate system allows tracking of stochastic cell populations, and importantly integrating data from different samples, thus enabling multimodal integration of data that do not necessarily need to be *a priori* spatially resolved. We therefore propose to use our coordinate system for integrated spatial transcriptomics (Karaiskos et al., 2017; Maseda et al., 2021; Satija et al., 2015), hopefully establishing the full nature of endothelial heterogeneity that drives vascular patterning.

## Methods

### Cdc42 and Rac1 endothelial knock-out

The following mice strains were used: Vegfr3-CreERT2;mTmG (Martinez-Corral et al., 2016; Muzumdar et al., 2007; Tammela et al., 2011), *Ccd42^flox/flox^*(Wu et al., 2006) and *Rac1^flox/flox^* (Chrostek et al., 2006). Mice were maintained at the Max Delbruck Center for Molecular Medicine under standard husbandry conditions. All experiments were approved by local authorities (LAGeSo, animal licence G0207/18, 14/11/2018). Both male and female mice were used. To induce Cre-mediated recombination, pups were injected once intraperitoneally with (Z)-4-hydroxytamoxifen hydroxytamoxifen (Sigma-Aldrich 7904, 10 µg/g of 1 mg/ml peanut oil solution) at postnatal day 5 (P5) or P8. Eyes were collected at P6, P7, P8 or P9. For loss-of-function experiments, Vegfr3-CreERT2;mTmG were crossed with *Ccd42^flox/flox^* and *Rac1^flox/flox^*. Gene deletion in ECs was observed with the expression of GFP as a proxy for Cre-flox function in combination with genotyping.

### Retina preparation and image acquisition

Retinas were dissected, flattened, stained and fixed as described in (Neto et al., 2018). To label EC membranes and EC nuclei, retinas were stained for Isolectin 4 (IB4, Invitrogen) and (ETS)-related gene (ERG, Abcam). Imaging was performed on a laser scanning confocal fluorescence microscope (Carl Zeiss LSM700, LSM780). Three channels were acquired: (1) GFP-signal of the ECs in which Cre-recombination occurred, (2) ERG signal of all EC nuclei, and (3) IB4 signal of the entire vasculature. To normalize for EC size in the distribution analysis, the GFP image was multiplied with the ERG image.

### Image Analysis of *in vivo* data

The IB4 channel (after maximum projection) was used to manually label veins, arteries and the optic nerve in Fiji (Schindelin et al., 2012). For every pixel in the image, three numbers were computed (using the masks as referential): (1) distance to the nearest vein (*d*_v_); (2) distance to the nearest artery (*d*_a_); and (3) radial distance to the optic nerve (*d_r_*). From these measures, the relative distances by ϕ_v-a,rp_ = *d*_v_/(*d*_v_ + *d*_a_) were obtained. Furthermore, we labelled straight line extensions of vein and arteries into the sprouting front. In a similar fashion we computed a modified vein-artery ϕ_v-a,sf_ as a reference system in the sprouting front from ϕ_v-a,sf_ = d_v,sf_/(d_v,sf_ + d_a,sf_), where d_v,sf_ and d_a,sf_ denote the distances to the nearest straight line extension of a vein and artery, respectively We distinguished the remodelling plexus and the sprouting front by adding an elliptical mask that approximately marks the part of the retina already containing mature vessels (see Box Figure 1). The EC distribution was computed by performing the operation for 10000 randomly drawn GFP and ERG-positive pixels in each retina, which were used as a proxy for EC distribution. A kernel density estimation was used to approximate the underlying EC distribution in the two-dimensional coordinate system spanned by ϕ_v-a_ and *d_r_*. For computational analysis, a Python-based workflow was developed, accessible on GitHub (https://github.com/wgiese/EC_migration_retina).

### Kernel density estimation (KDE)

A kernel density estimation (KDE) is used to approximate the underlying probability density function that is generated by counting positively labelled pixels in the coordinate system spanned by ϕ_v-a,rp_ and r, and ϕ_v-a,sf_ and r, respectively. In contrast to using discrete bins, the KDE plot is based on an approximation by Gaussian kernels, which results in a continuous density estimate. Scotts method (Scott, 1979) was used to estimate the Gaussian kernel bandwidth. For visualization, we employed the seaborn package from python.

### Cell culture and siRNA treatment

Commercialized human umbilical vein endothelial cells (HUVECs, pooled donors, PromoCell) were mycoplasma tested and routinely cultured following the manufacturer’s guidelines. Cells were used at passages 3-5 for experiments and cultured at 37°C and 5% CO_2_ in complete EC basal medium containing growth factors EGM2-Bulletkit (Lonza) to ensure a stable environment for optimal cell growth. During every cell passage or during seeding in mechanical assay chambers, cells were washed once in sterile PBS, then incubated for 5min in trypsin/EDTA at 37°C, 5% CO_2_, and quenched with EGM2 and FBS. Following centrifugation and cell count, cells were seeded for experiments or transfection treatment. For siRNA experiments, HUVECs were transfected with the corresponding SMARTpool: siGENOME negative control (Dharmacon), CDC42 (OnTARGETplus SMARTpool, L-005057-00-0005, Dharmacon) and RAC1 (OnTARGETplus SMARTpool, L-003560-00-0005, Dharmacon). Briefly, HUVECs were seeded at 60–70% confluence to a 6-well plate and incubated overnight in complete medium. At the day of transfection, cells were washed using sterile PBS and transfected with 50 µM siRNA using Dharmafect-1 (Dharmacon) transfection reagent following manufacturer protocols, in antibiotics free EGM2. Following 24h of incubation, the reagents were replaced with fresh complete medium. Knockdown efficiency was validated 48 h and 72 h post transfection using qPCR (see Figure 5 Supplement 2).

### Live imaging experiments and Cell tracking

#### Shear stress experiments

For live-imaging experiments, HUVECs were seeded onto 0.4 Luer slides (Ibidi) at a concentration of 2 million cells/ml, 28-30h post siRNA knockdown. The following day, nuclear dye SPY505 (Spyrochrome) was added with CO_2_ independent MV2 media (PromoCell) and incubated at 37°C for 4h prior to imaging. Nuclear tracking assays were carried out 52 hours after siRNA transfection and EC nuclei were tracked for 17 hours using a confocal microscope (Carl Zeiss, LSM 980), equipped with a Plan-Apochromat 20×/0.8 NA Ph2 air objective and the Definite Focus system, by imaging a 3×3 tile scan, 3 slice z levels per tile, with a temporal resolution of 5 minutes and a pixel resolution of 1,2068 pixels/μm, using a 488 excitation laser at low power (0.5%). Migration analysis was carried out on the stitched images using the TrackMate Fiji plugin with StarDist (Schmidt et al., 2018) for nuclei tracking. Track records obtained from this analysis were used to create individual and pooled trajectory plots and migration velocity plots (directional and general) as shown in Figure 5, using a Python script (available at https://github.com/wgiese/ec-tracking-analysis).

#### Scratch wound assay

For live-imaging experiments, HUVECs were incubated with Hoechst 33342 (Thermo Fisher Scientific) for 20min at 37°C before transferring samples for imaging under controlled temperature and CO_2_ levels. For wound assay experiments, cultured cells were seeded in Culture-Insert 2-Well (Ibidi). Time-lapse imaging was performed on a confocal microscope (Carl Zeiss, LSM 780), equipped with a Plan-Apochromat 20×/0.8 NA Ph2 air objective and the Definite Focus system. Time series of single planes were acquired sequentially from randomly selected positions every 15 min at a resolution of 1,2044 pixels/μm using a 405 nm excitation laser at low-power and a transmitted light detector (DIC). Segmentations, cleanings, and trimming of individual nuclei tracks were obtained using Fiji and CellProfiler. In wound assay monolayers, a mask was applied to only track cells localized within the initial monolayer’s leading front (200 μm). Briefly, to monitor the moving trace of each nucleus, the nuclei identified at each time frame were linked based on the nearest neighbour method. In case of mitotic events, one of the daughter cells was directly linked to the mother cell, the other was assigned a new number, and its correlation with the mother cell was saved. Quantification was obtained only from tracks accumulating a minimal number of consecutive frames. Accumulated distance represented the sum of individual displacements of the cell over each interval of the cell track, 15 min per frame. Cell velocity was determined based on accumulated displacements divided by the total tracked time. Euclidean distance or total displacement was defined as the straight-line distance between the initial and final position of a tracked cell. Directness characterises the straightness of cell trajectories. Additionally, the forward migration index represents the efficiency of the forward migration of cells and how they relate to the direction in which the entire cell population drifts. Visualizations and data extraction were obtained using the Chemotaxis and Migration Tool (Ibidi).

### Computational Model

We constructed a computational model to estimate EC migration speed and asses the strength of the two competing shear stress and VEGFA-attractor cues in the remodelling plexus and sprouting front. The movement of ECs from one lattice site to another is described by a step selection function 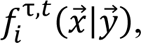 which describes the probability of an EC agent *i* moving from position 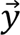 at time *t* to position 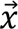 at time *t* + *τ*. Here, 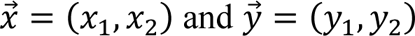 are two-dimensional vectors. The step selection function comprises four different aspects of cellular movement, which are (i) biased Brownian movement, (ii) signalling cues, (iii) mechanical cues and (iv) the cellular state. The computational model is based on a Gillespie algorithm (Gillespie, 2007), the model framework has recently been adapted to EC movement on a spatial grid by (Stepanova et al., 2021). Given that an EC agent *i* is at site 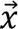, the probability distribution for an agent moving from site 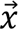 to site 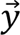 is given by the following step-selection function:

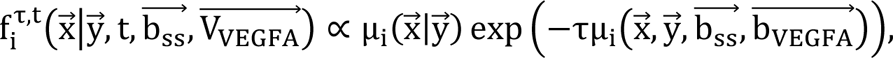

which is dependent on local cues in the vascular bed and is described by two vector fields: (i) the shear stress field 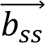 and (ii) the VEGFA induced signalling cue 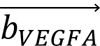. The VEGFA gradient is modelled according to (Claxton and Fruttiger, 2005). The simulation with several hundred EC agents could be performed on any defined patch of our annotated retinal images.. The code is available on GitHub https://github.com/wgiese/ec-retina-migration-model.

### Parameter estimation

We ran 250 Simulations each to construct two phase diagrams showing the dependency of (i) coupling strength and movement rate as well as (ii) coupling strength and proportion of the shear stress cue. Next, we performed a Gaussian process regression by using the *gaussion_process* module from scikit-learn (Pedregosa et al., 2011). The standardised mean squared error (SMSE) was used as error metric meaning (x_sim_ − x_exp_)^2^/s^2^ where x_sim_ is the sample mean of the simulated quantity, xexp is the sample mean of the experimentally measured quantity and s is the standard deviation of the experimentally measured quantities. The SMSE was computed for the 10^th^ percentile, median and 90^th^ percentile at every time point for the vein-to-artery distances in the remodelling plexus and sprouting front as well as the distances to the optic nerve.

## Statistical analysis

All statistical analysis was performed using Python packages *Statannotations* (Charlier et al., 2022), *scipy.stats* and *Seaborn*. Statistical details of experiments are reported in the figures and figure legends. Sample size is reported in the figure legends and no statistical test was used to determine sample size. The biological replicate is defined as the number of cells, images, animals, as stated in the figure legends. No inclusion/exclusion or randomization criteria were used, and all analysed samples are included. Comparisons between two experimental groups were analysed using Welch’s t-test. We considered a result significant when p<0.05.

## Acknowledgments

We thank Marie Altmann and all the animal caretakers at the Max Delbrück Center for Molecular Medicine and, furthermore, Alexandra Klaus-Bergmann for her help with advice and application writing. We would also like to thank Alexandre Dubrac and the entire AG Gerhardt for helpful discussions.

## Author contributions

**Wolfgang Giese:** Conceptualization; data curation; formal analysis; investigation; methodology; software; project administration; validation; visualization; writing – original draft; writing – review and editing. **André Rosa:** Conceptualization; data curation; formal analysis; investigation; methodology; project administration; validation. **Elisabeth Baumann:** Data curation; formal analysis; investigation; methodology; validation; visualization; writing – review and editing. **Olya Oppenheim:** Data curation; investigation; validation; visualization; writing – review and editing. **Emir B. Akmeric:** Data curation; investigation; validation; visualization; writing – review and editing. **Santiago Andrade:** Data curation; investigation; validation; visualization; writing – review and editing. **Irene Hollfinger:** Investigation. **Eireen Bartels-Klein**: Investigation. **Silvanus Alt:** Conceptualization, methodology. **Holger Gerhardt:** Conceptualization; funding acquisition; methodology; project administration; supervision; writing – review and editing.

## Supplementary Figures

**Figure 1 Supplement 1:**
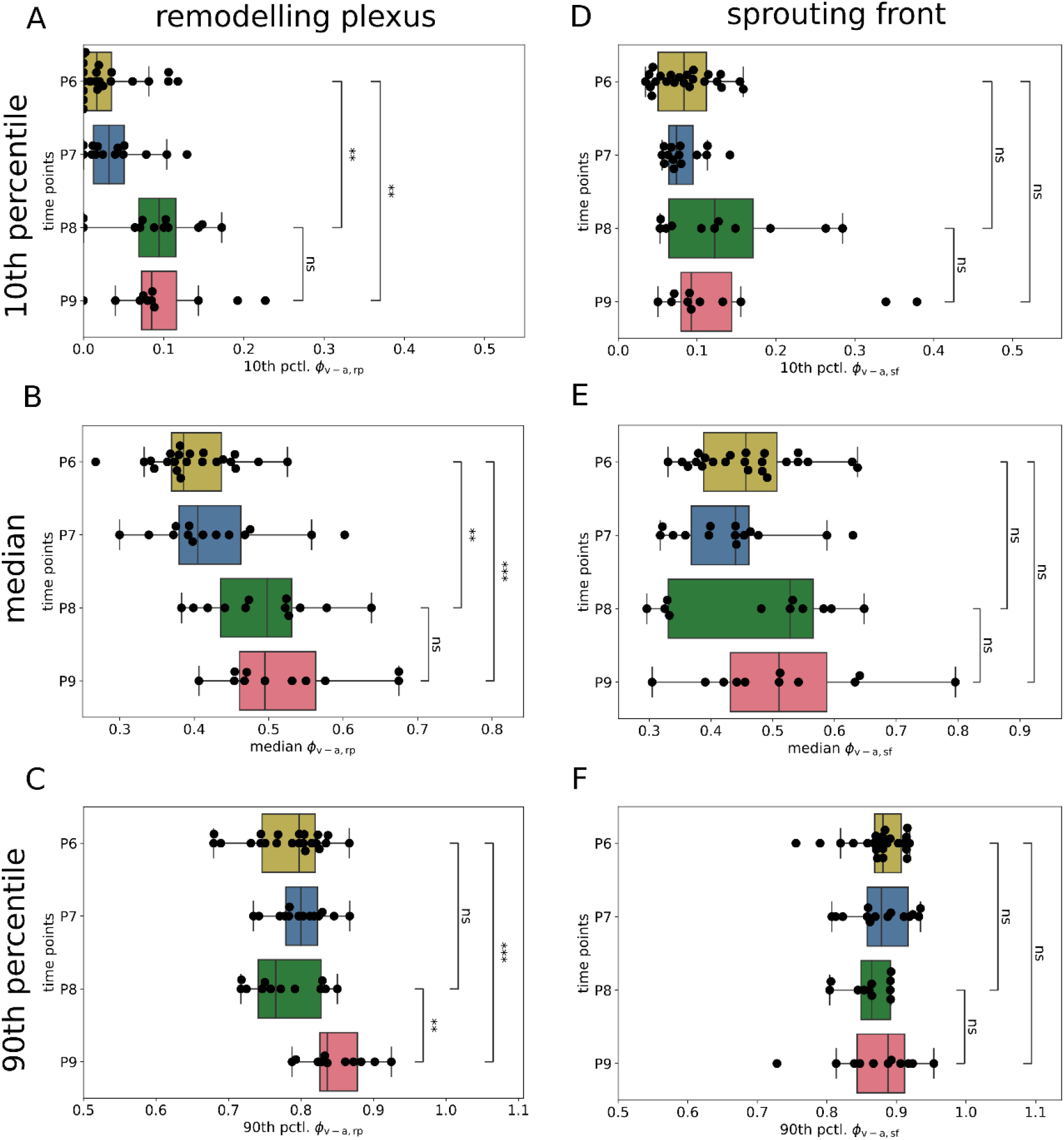
Temporal evolution of the 10th, 50th and 90th percentile of the EC distribution in the remodelling plexus and sprouting front in control retinas. For each mouse retina sample (indicated by a black dot) we computed the 10th, 50th (median) and 90th percentile of the EC distribution for the remodelling plexus and sprouting front along the ϕ_v-a,rp_ and ϕ_v-a,sf_ coordinate, respectively. **A**: In the remodelling region, we observed a significant shift of the 10^th^ percentile of the labelled EC population towards arteries over time comparing P6 to P8 (p < 0.01) and P6 to P9 (p < 0.01), but not for the last time interval comparing P8 to P9. **B:** Similarly, we observed a significant shift of the median of the labelled EC population towards arteries over time comparing P6 to P8 (p < 0.01) and P6 to P9 (p < 0.001), but again not for the last time interval comparing P8 to P9. **C:** For the 90^th^ percentile, the population shift in the remodelling plexus was delayed with significant difference comparing P6 to P9 (p < 0.001) and P8 to P9 (p < 0.01), but not for the last time interval comparing P6 to P8. **D, E, F**: In the sprouting front, we observed a slight but not significant progression from P6 to P8 and P6 to P9 for the median, and none for the 10th and 90th percentile.

**Figure 1 Supplement 2:**
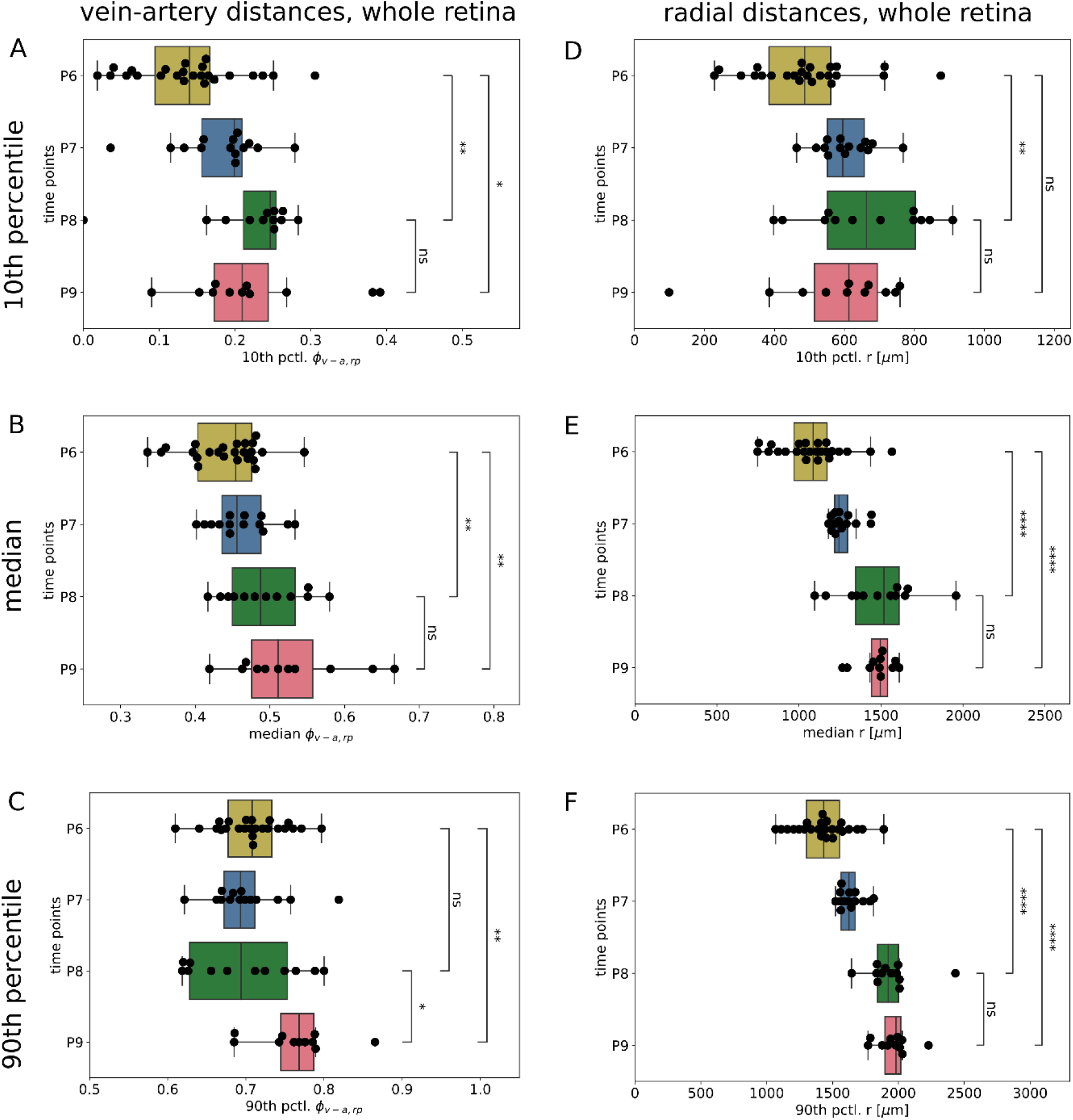
Temporal evolution of the 10th, 50th and 90th percentile of the entire EC distribution in control retinas. For each mouse retina sample (indicated by a black dot) we computed the 10th, 50th (median) and 90th percentile of the EC distribution in the whole retina in the (ϕ_v-a,rp_, r)-coordinate system. **A, B**: For the 10^th^ percentile and median, we observed significant differences in the distribution along ϕ_v-a,rp_ comparing P6 and P8 as well as P6 and P9, but did not observe any significant differences comparing P8 to P9. **C**: For the 90^th^ percentile, the population shift towards the artery occurred with a delay, meaning that there was a significant difference between point P6 and P9 as well as P8 and P9, but not from P6 and P8. **D, E, F**: For the EC progression in radial direction, we observed a significant shift from P6 to P8 for 10th, 50th (median) and 90th percentile, while the progression from P8 to P9 was stagnant. The differences comparing P6 and P9 were significant for the median and 90^th^ percentile, but not for the 10^th^ percentile.

**Figure 2 Supplement 1:**
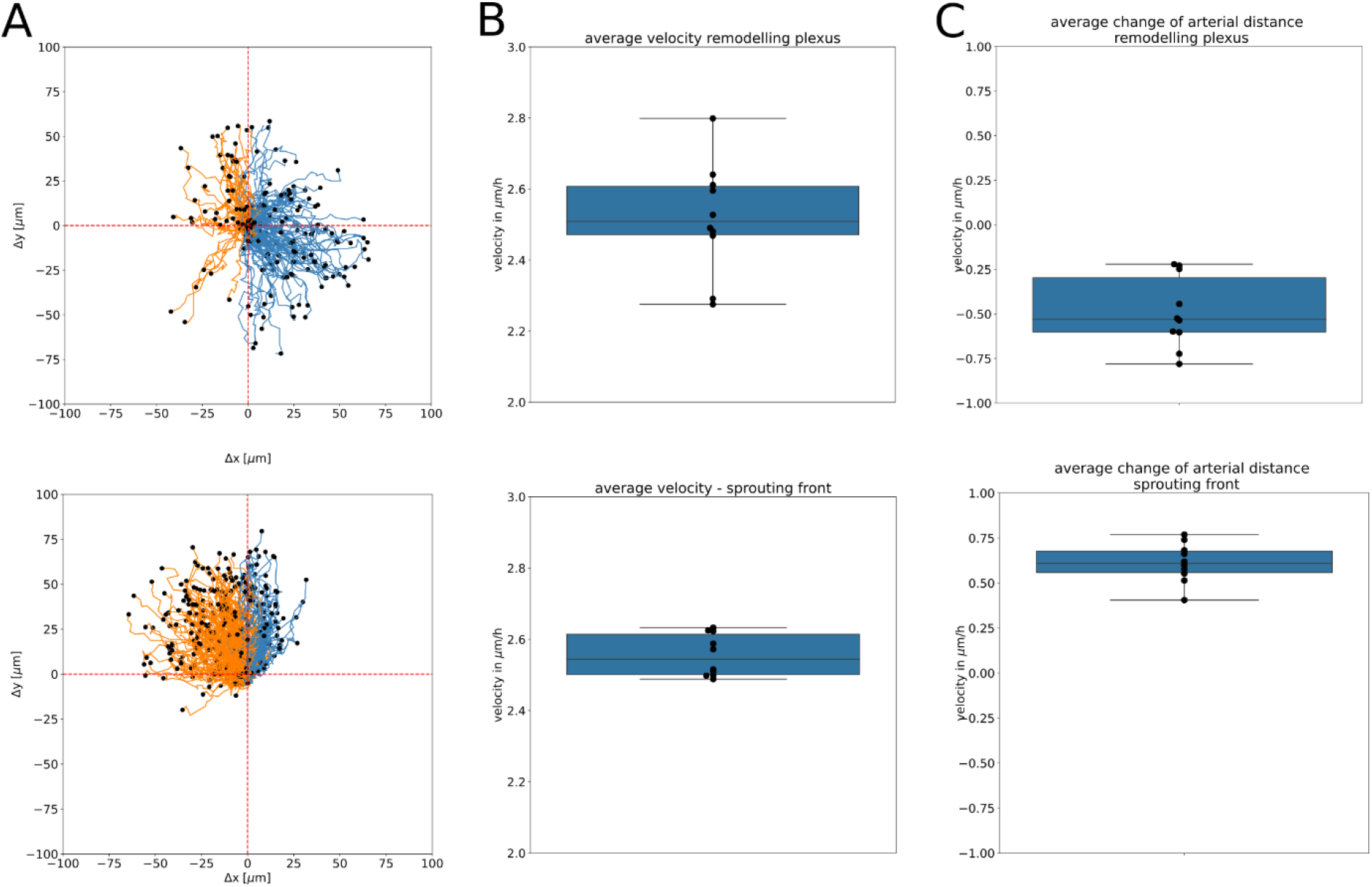
EC trajectory plots and velocity estimates. **A**: Trajectory plots of n = 500 simulated ECs over an observational time period of one day from P6 to P7 in the remodelling plexus and sprouting front. **B:** Total velocity estimates in the remodelling plexus and sprouting front. **C:** Velocity in arterial direction that is computed from the distance change of Δ*d*_*a*_ over time in the remodelling plexus and from Δ*d*_*a*,*sf*_ over time in the sprouting front. A positive velocity indicates movement away from the artery, while a negative velocity indicates movement towards the artery.

**Figure 3 Supplement 1:**
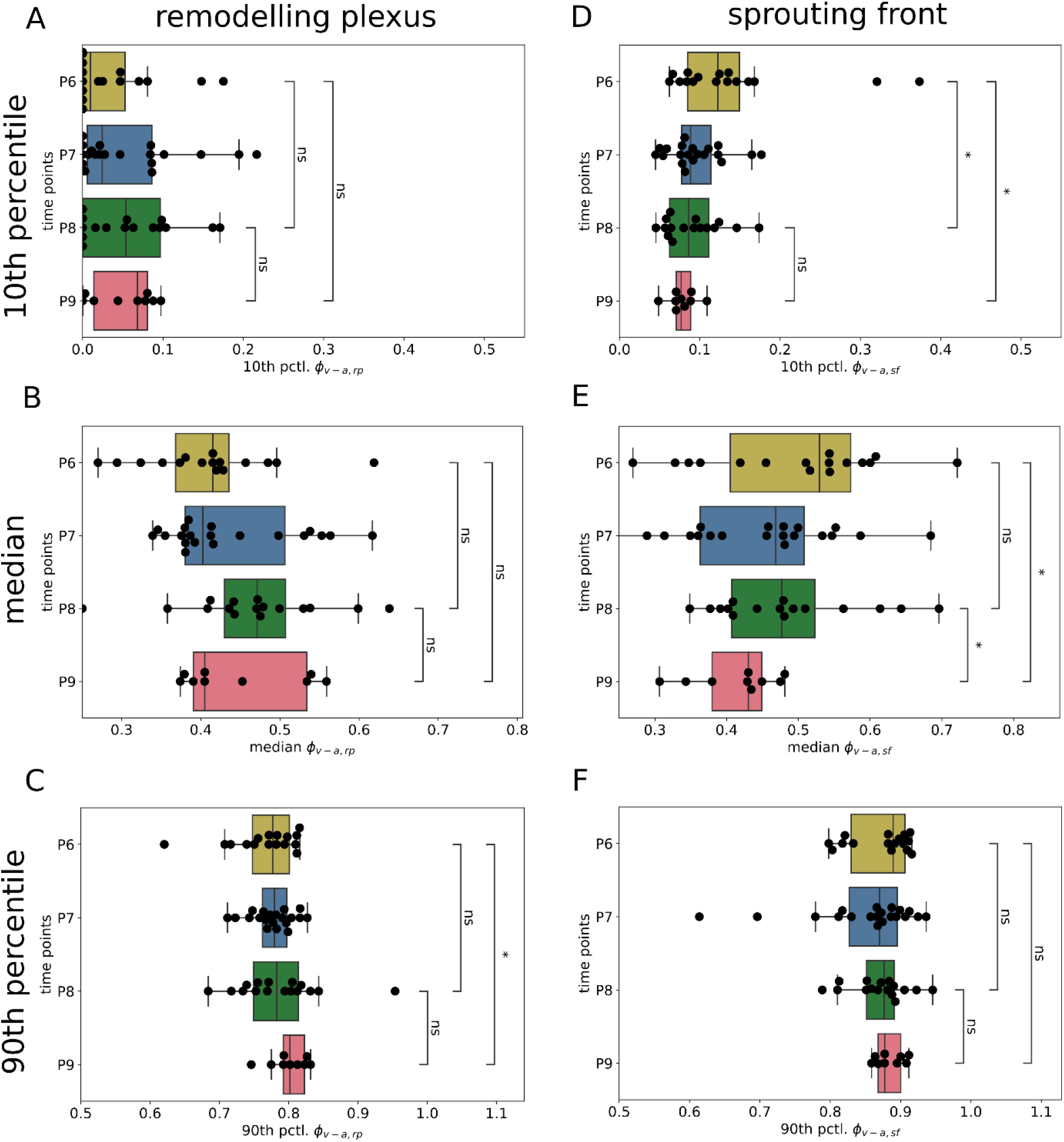
Temporal evolution of the 10^th^, 50^th^ and 90^th^ percentile of the Cdc42 depleted EC population in the remodelling region and sprouting front. For each mouse retina sample (indicated by a black dot), we computed the 10th percentile, 50th (median) and 90th percentile of the EC distribution for the remodelling region and sprouting front along the ϕ_v-a,rp_ and ϕ_v-a,sf_ coordinate, respectively. **A, B**:. We did not observe any significant population shift for the 10th percentile and median in the remodelling region. **C**: For the 90th percentile we observed a significant shift of the population towards arteries comparing P6 and P9, but not did not observe significant differences comparing P6 to P8 and P8 to P9. **D**: In contrast, in the sprouting front we observed a significant population shift towards veins for the 10th percentile comparing P6 and P8 and P6 to P9. **E**: Similarly, but with a delay, we observed a shift of the median towards veins in the sprouting front comparing P6 to P9 and P8 to P9. **F**: For the 90^th^ percentile, we did not observe any significant differences in the sprouting front. Differences between time point P6 and P8, P6 and P9 as well as P8 and P9 were assessed statistically by Welch’s t-test.

**Figure 3 Supplement 2:**
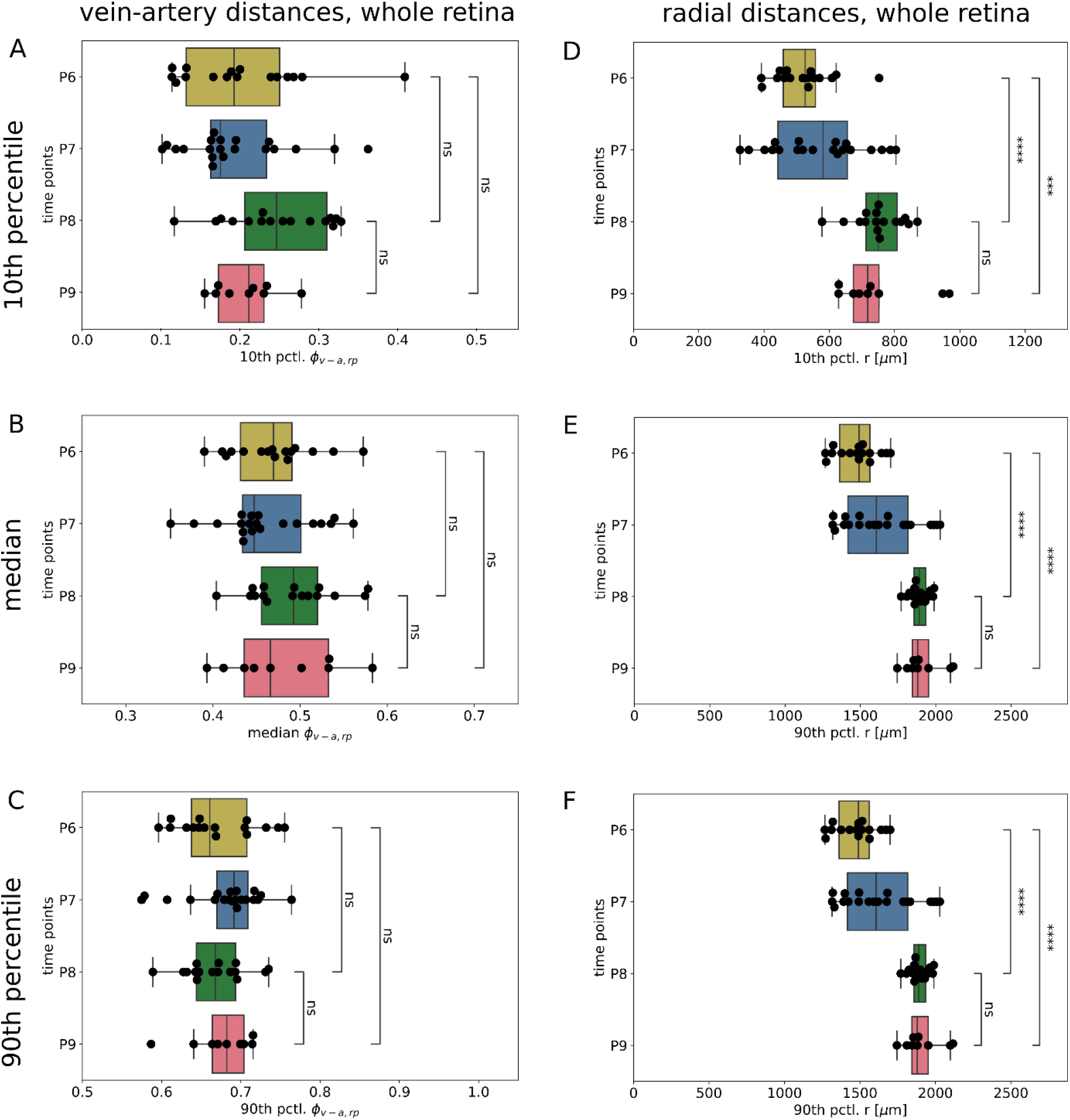
Temporal evolution of the 10th, 50th and 90th percentile of the Cdc42 depleted EC population in the whole retina. For each mouse retina sample (indicated by a black dot) we computed the 10th, 50th (median) and 90th percentile of the EC distribution for the whole retina in the (ϕ_v-a,rp,_r)-coordinate system. Differences between time point P6 and P8, P6 and P9 as well as P8 and P9 were assessed statistically by Welch’s t-test. We did not observe any significant population shift in ϕ_v-a,rp_ for the 10th and 90th percentile, while for the median a significant shift in ϕ_v-a,rp_ towards veins was observed from P8 to P9. For the EC progression in the radial direction, we observed a significant shift from P6 to P8 for the 10th, 50th (median) and 90th percentile, while the progression from P8 to P9 was stagnant.

**Figure 3 Supplement 3:**
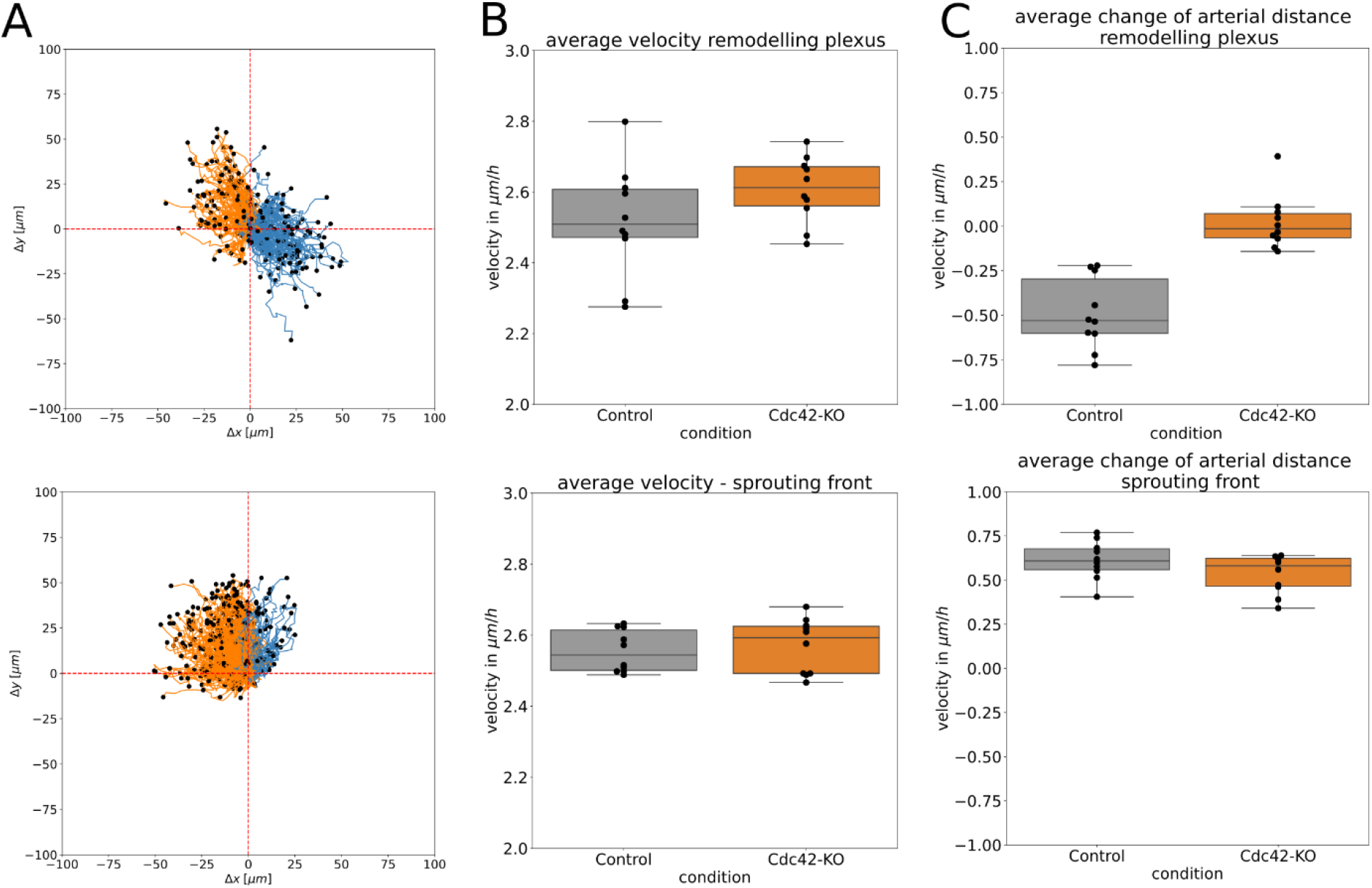
EC trajectory plots and velocity estimates for Cdc42 depleted ECs. **A**: Trajectory plots of n = 500 simulated ECs over an observational time period of one day from P7 to P8 in the remodelling plexus and sprouting front. **B:** Total velocity estimates in the remodelling plexus and sprouting front of Cdc42 depleted cells in comparison to the control. **C:** Velocity in arterial direction that is computed from the distance change of Δ*d*_*a*_ over time in the remodelling plexus and from Δ*d*_*a*,*sf*_ over time in the sprouting front of Cdc42 depleted cells in comparison to the control. A positive velocity indicates movement away from the artery, while a negative velocity indicates movement towards the artery.

**Figure 4 Supplement 1:**
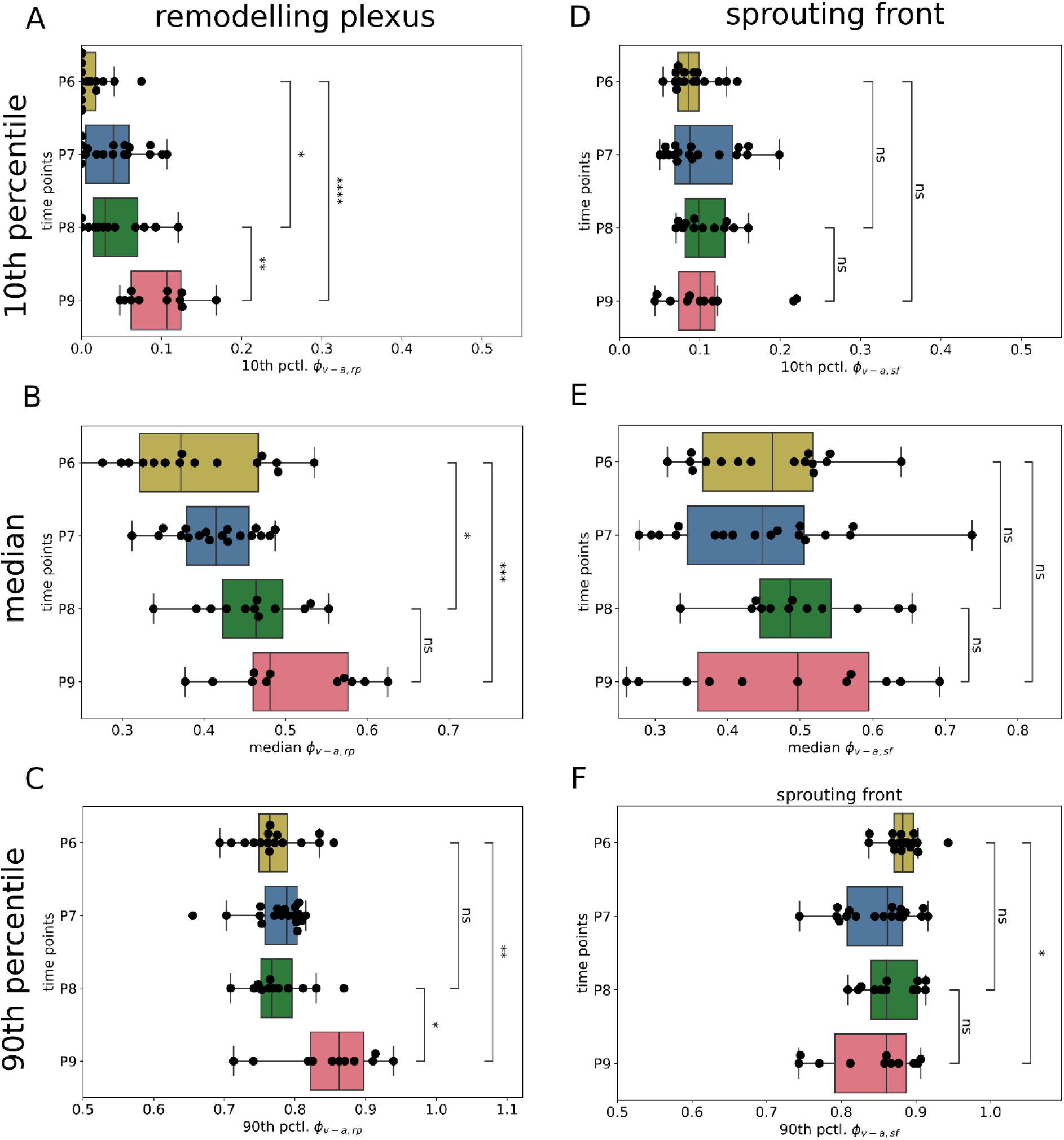
Temporal evolution of 10th, 50th and 90th percentile of the EC distribution deficient in Rac1 in the remodelling region and sprouting front. For each mouse retina sample (indicated by a black dot), we computed the 10th, 50th (median) and 90th percentile of the EC distribution for the remodelling region and sprouting front along the ϕ_v-a,rp_ and ϕ_v-a,sf_ coordinate, respectively. **A, B, C**: The progression of the 90th percentile occurred with a delay in the remodelling plexus, meaning that the shift from P6 to P8 was not significant, while we observed significant progression from P8 to P9 as well as from P6 to P9. **D, E**: In the sprouting front, we did not observe any significant differences for the 10^th^ percentile and the median distribution, however, we observed a significant change of the 90th percentile from P6 to P9 backwards in the direction of veins. Differences between time point P6 and P8, P6 and P9 as well as P8 and P9 were assessed statistically by Welch’s t-test.

**Figure 4 Supplement 2:**
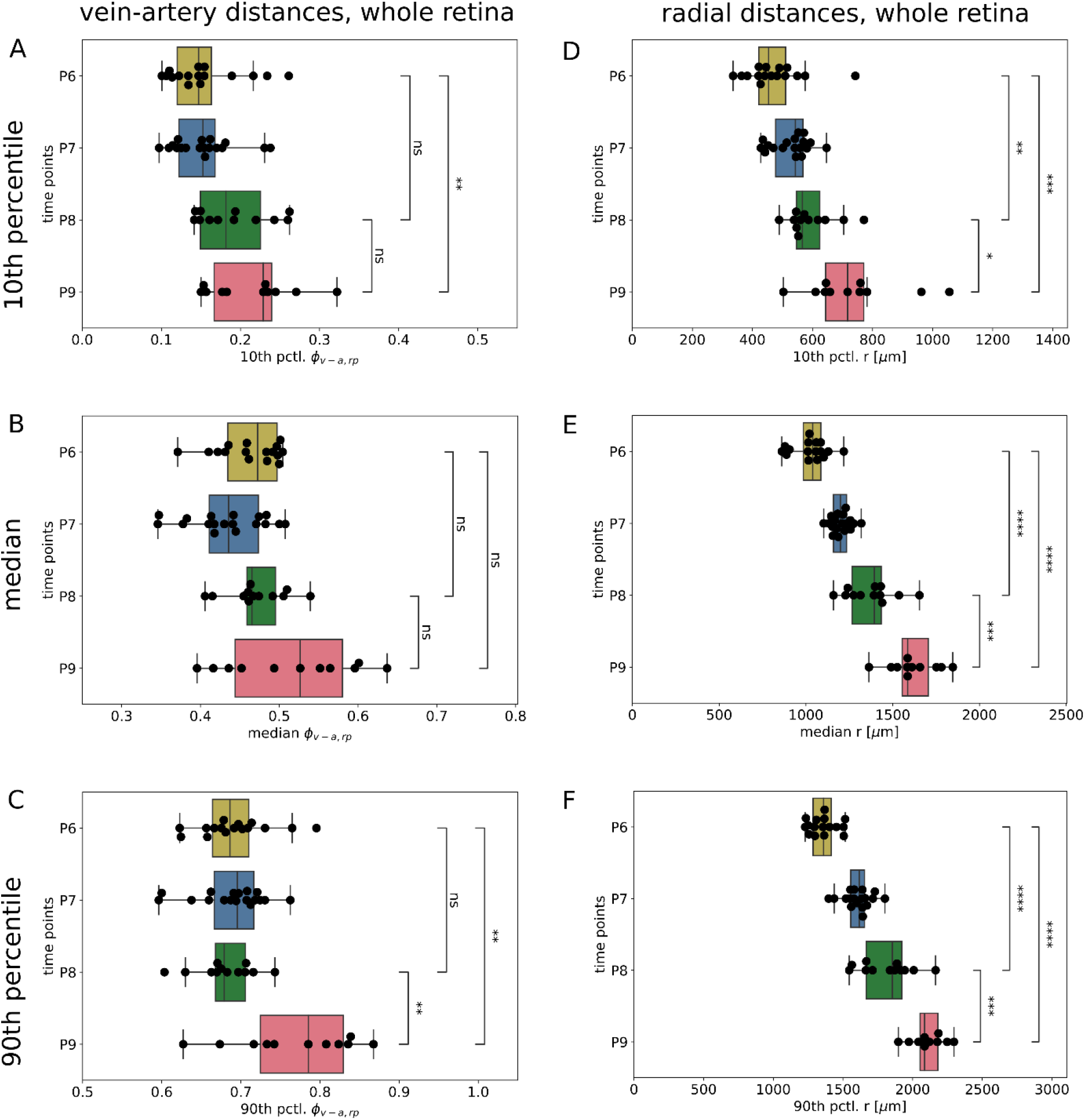
Temporal evolution of the 10^th^, 50^th^ and 90^th^ percentile of the Rac1 depleted EC population in the whole retina. For each mouse retina sample (indicated by a black dot), we computed the 10th, 50th (median) and 90th percentile of the EC distribution for the remodelling region and sprouting front in the (ϕ_v-a,rp,_r)-coordinate system. Differences between time point P6 and P8, P6 and P9 as well as P8 and P9 were assessed statistically by Welch’s t-test. **A, B, C**: A significant shift of the population towards arteries is observed for the 10^th^ and 90^th^ percentile, but not for the median. We see a significant population shift in ϕ_v-a,rp_ direction for the median and 90th percentile from P6 to P9. **D, E, F**: For the EC progression in radial direction, we observed a significant shift from P6 to P8, P6 to P9 and P8 to P9 for the 10th, 50th (median) and 90th percentiles.

**Figure 4 Supplement 3:**
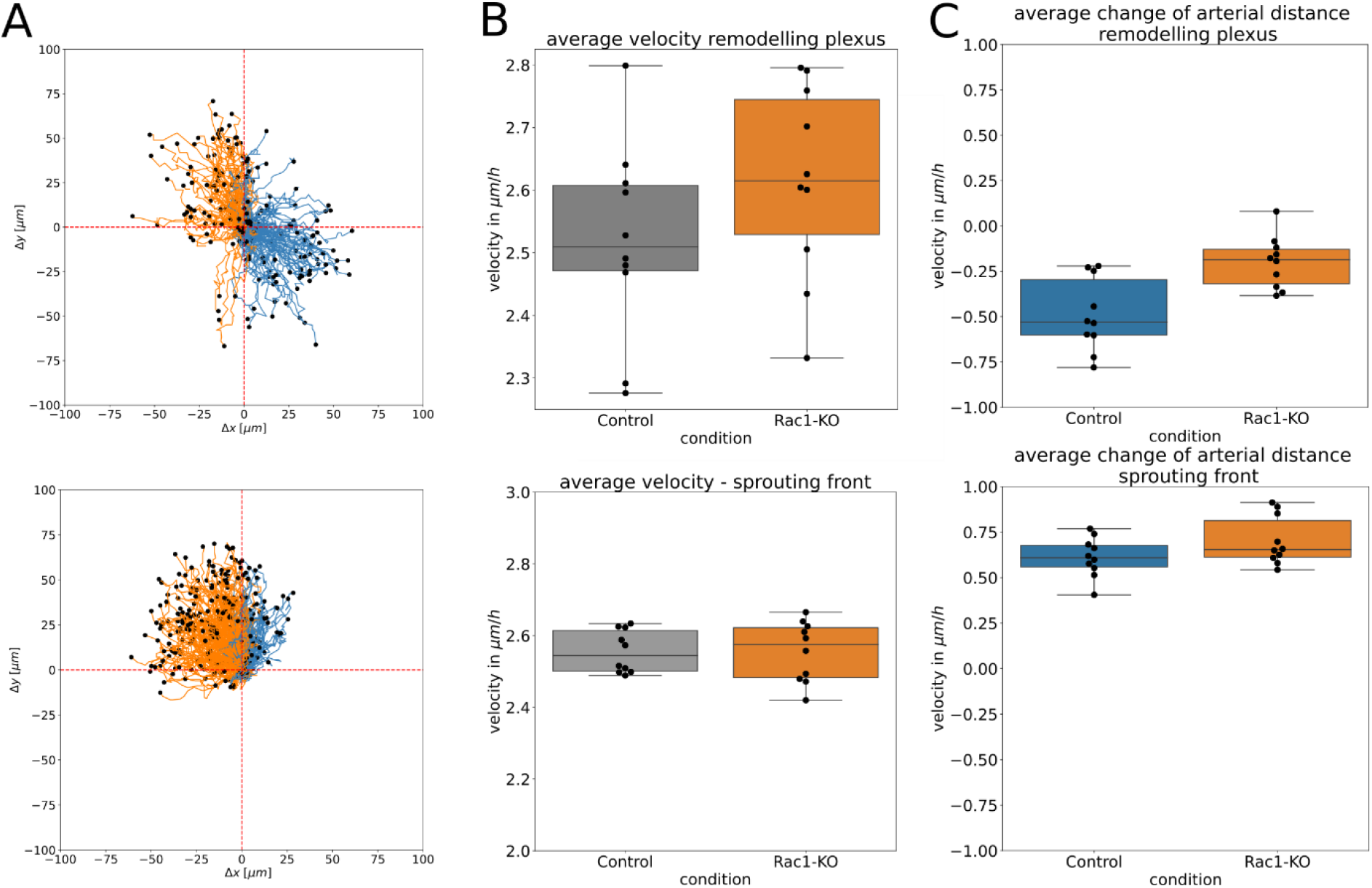
EC trajectory plots and velocity estimates for Rac1 depleted ECs. **A**: Trajectory plots of n = 500 simulated ECs over an observational time period of one day from P7 to P8 in the remodelling plexus and sprouting front. **B:** Total velocity estimates in the remodelling plexus and sprouting front of Rac1 depleted cells in comparison to the control. **C:** Velocity in arterial direction that is computed from the distance change of Δ*d*_*a*_ over time in the remodelling plexus and from Δ*d*_*a*,*sf*_ over time in the sprouting front of Rac1 depleted cells in comparison to the control. A positive velocity indicates movement away from the artery, while a negative velocity indicates movement towards the artery.

**Figure 5 Supplement 1:**
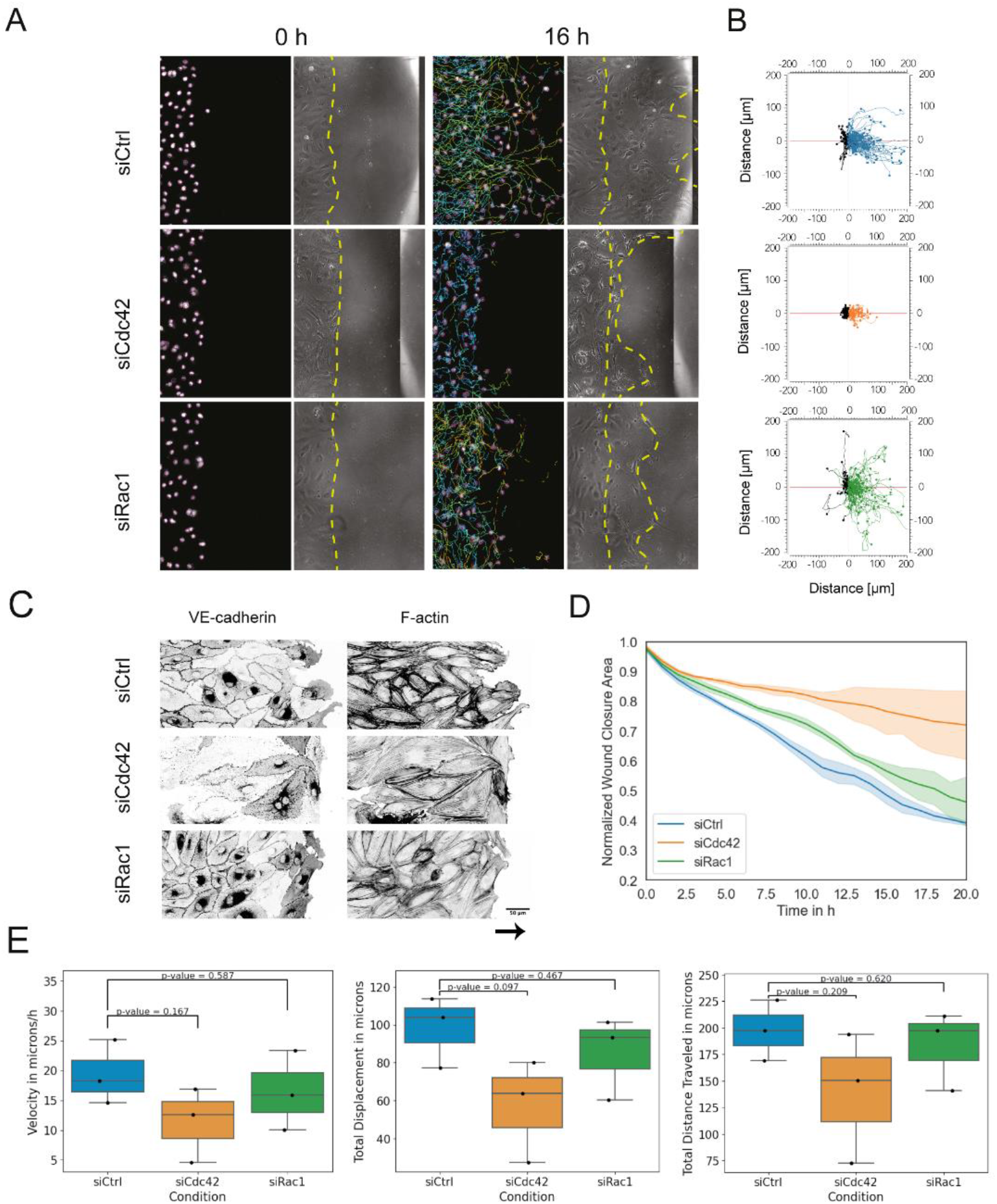
Impact of Cdc42 and Rac1 depletion on endothelial directional migration. Wound healing live-imaging assays: **A:** Representative images corresponding to siScr, siCdc42, and siRac1 conditions comparing initial and final time points (0 – 20h). Superimposed onto these images are EC trajectories, each shown in a different color. **B:** Migration plot of trajectories corresponding to leading-edge cells (within < 200 microns from the wound edge) in a representative region-of-interest per condition. Trajectories were superimposed and initiate from a zero point, indicating the direction of migration following the formation of a wound. Straighter directional migration is noticeable in siCtrl cells, while reduced migration is observed in siCdc42 cells. **C:** Immunostaining displaying VE-cadherin and F-actin in siCtrl, siCdc42, and siRac1migrating monolayers after 8 h of starting the migration assay. **D:** Kinetic analysis of trajectories for each condition. The data is normalized against the initial time point and represents the gap area between cell groups during a 20-hour period. Negative slopes enable comparison among conditions: the less negative the slope in the curve, the slower the migration. The slopes for each condition are as follows: siCtrl: −2.4642e-3, siCdc42: −0.7802e-3, and siRac1:-1.9996e-3. Data points represent biological replicates (n=3). **E:** Box plots depicting the total velocity in microns (left), total displacement in microns (center: measuring the straight-line distance between the initial and final position), and total distance travelled in microns (right: quantifying the cumulative path length travelled from the initial to final position) of trajectories for each condition. The data points correspond to biological replicates and are derived from trajectory means from the same experimental setup (n=3).

**Figure 5 Supplement 2:**
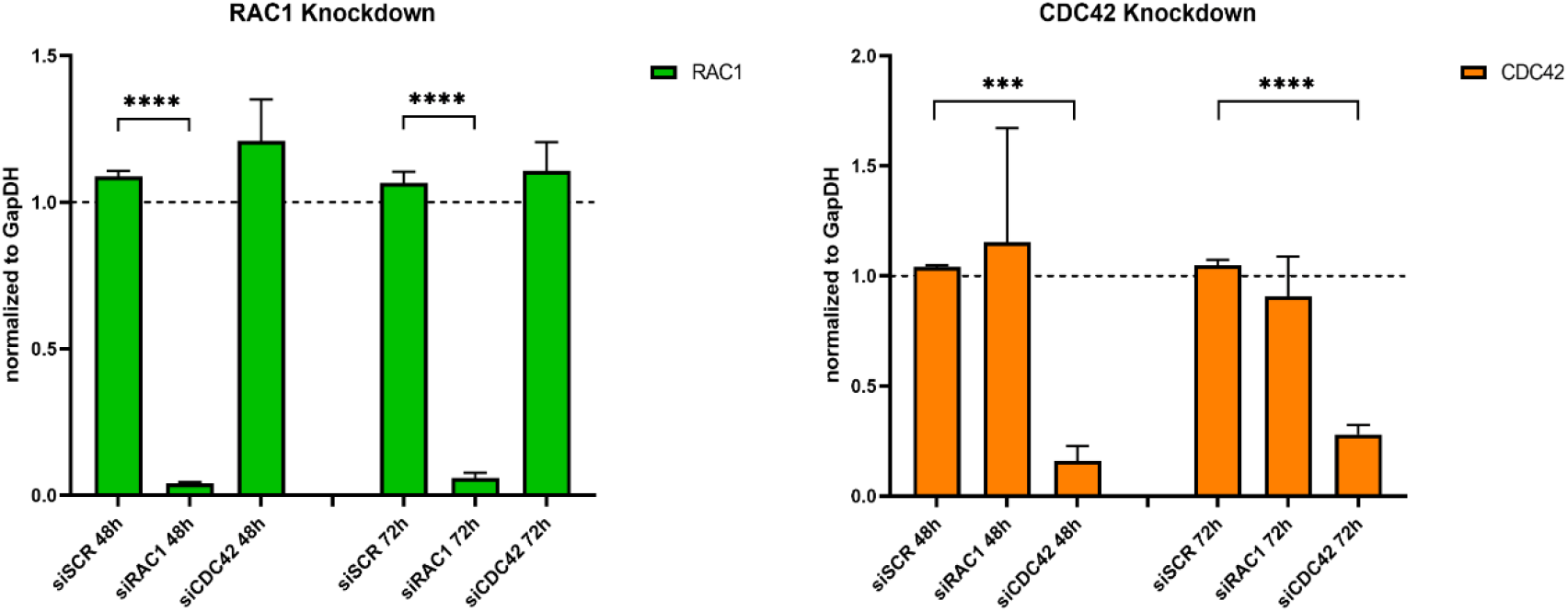
qPCR knockdown validation. Normalized expression ratio (N.E.R) of RAC1 (green) and CDC42 (orange) transcripts, 48h and 72h post siRNA transfection. Values are normalized to GapDH expression. Data points indicate the mean and standard error of n=3 for 48h and n=4 for 72h. Knockdown efficiency was statistically assessed by Welch’s t-test.

**Supplementary Figure 1:**
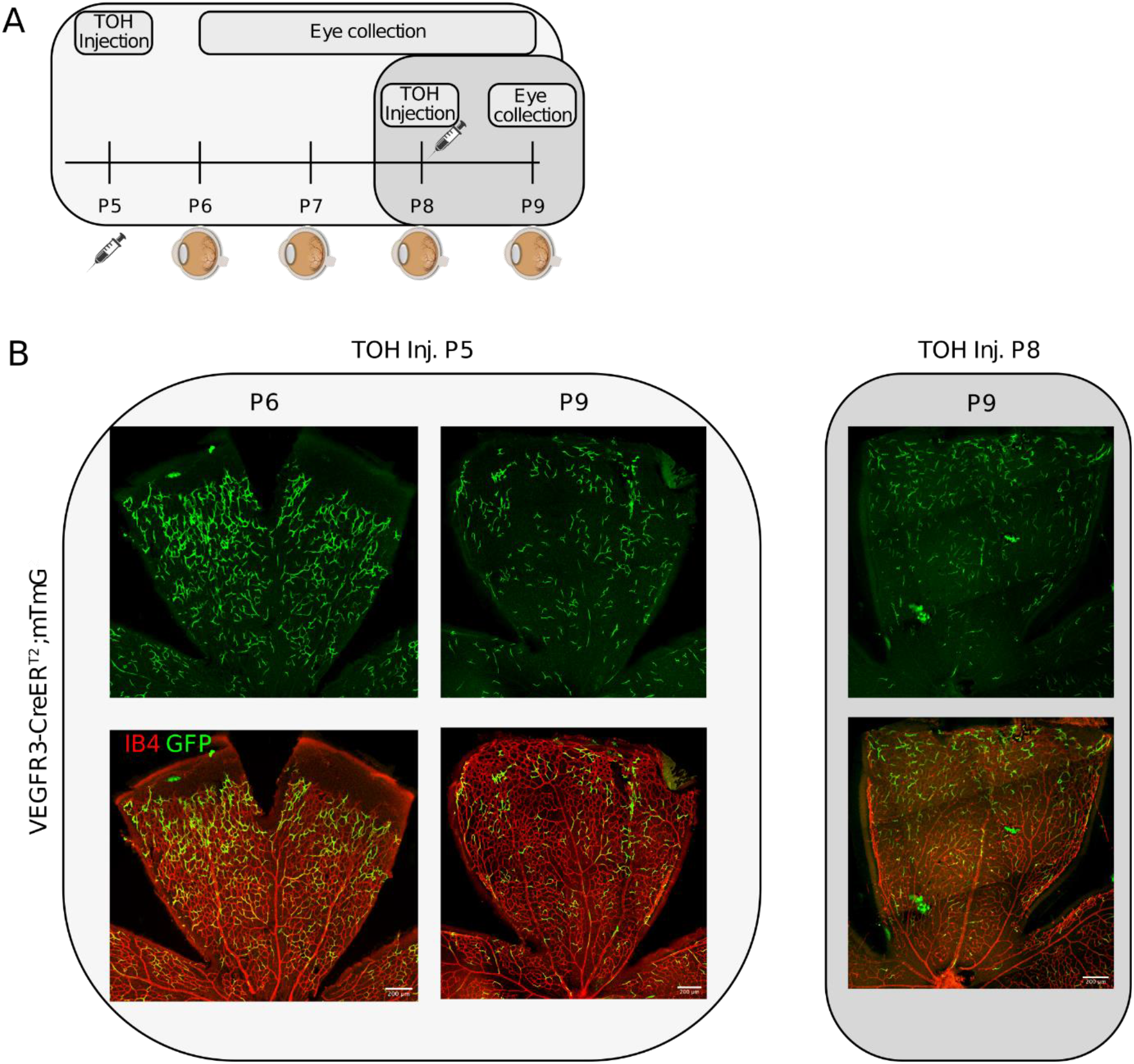
Tamoxifen injection scheme and retina collection. **A**: Pups were injected with tamoxifen at P5. Eyes were collected and EC distribution analysis was performed at P6, P7, P8 and P9. A control group of pups was injected at P8 and eyes were collected at P9. **B**: Pups injected at P5 showed randomly distributed GFP-labelling of ECs in veins, the remodelling plexus and the sprouting front but not in arteries at P6. By P9, GFP-labelled ECs were also present in arteries. In contrast, pups injected at P8 did not show GFP-labelling in arteries at P9.

**Supplementary Figure 2:**
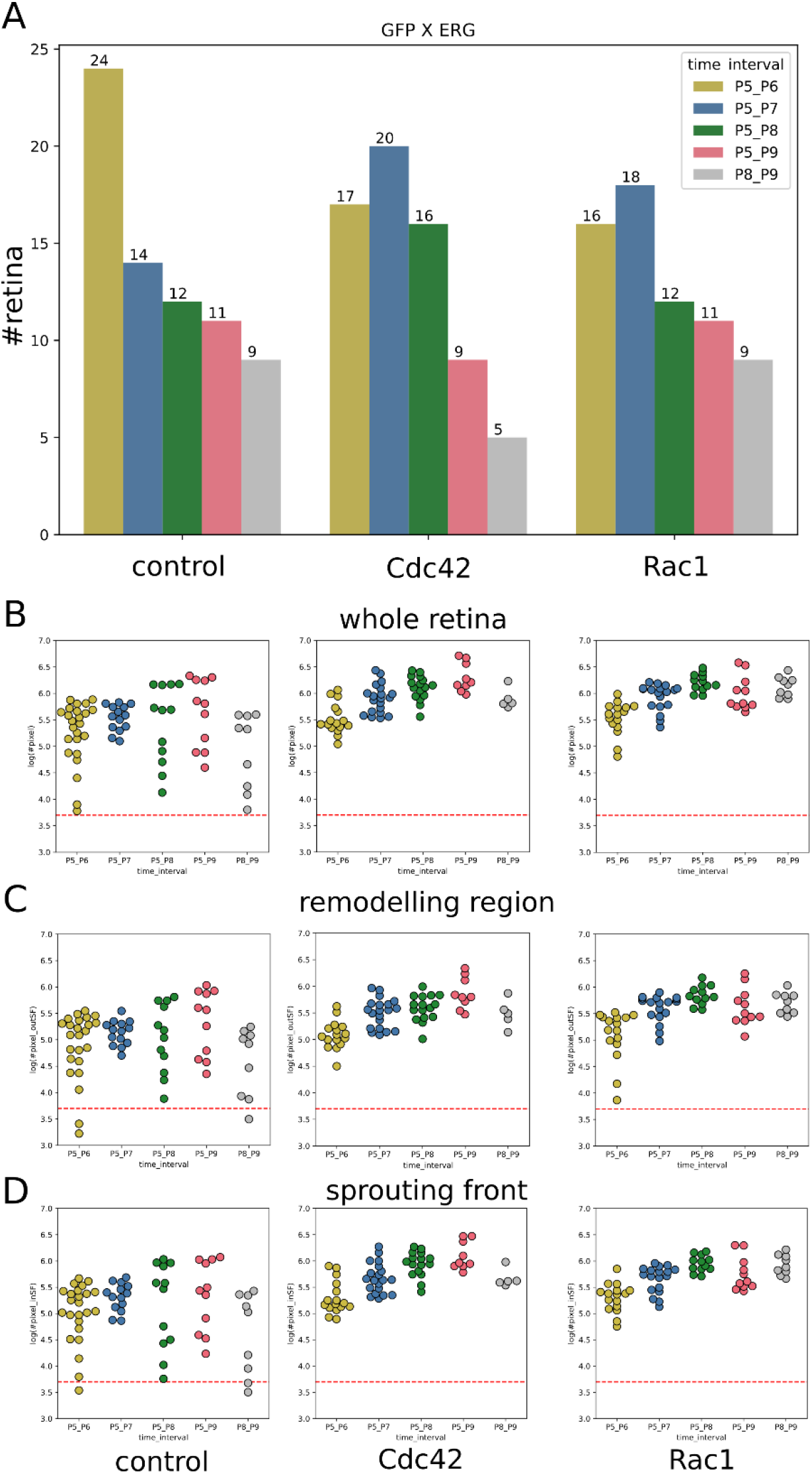
Numbers of mouse retina samples and GFPxERG positively labelled pixels in the remodelling region and sprouting front. **A**: Number of retina samples per time point and condition. **B, C** and **D**: Number of GFP and ERG positively labelled pixels in each retina microscopic image on a logarithmic scale. The minimum required number of pixels was set to 500 pixels and is indicated by dashed line in each plot. Microscopic images that do not meet that threshold were excluded from analysis. **B**: whole retina, **C**: only the remodelling region and **D**: only sprouting front.

## References

Ara T, Tokoyoda K, Okamoto R, Koni PA, Nagasawa T. 2005. The role of CXCL12 in the organ-specific process of artery formation. Blood 105:3155–3161. doi:10.1182/blood-2004-07-2563

Barbacena P, Dominguez-Cejudo M, Fonseca CG, Gómez-González M, Faure LM, Zarkada G, Pena A, Pezzarossa A, Ramalho D, Giarratano Y, Ouarné M, Barata D, Fortunato IC, Misikova LH, Mauldin I, Carvalho Y, Trepat X, Roca-Cusachs P, Eichmann A, Bernabeu MO, Franco CA. 2022. Competition for endothelial cell polarity drives vascular morphogenesis in the mouse retina. Dev Cell 57:2321–2333.e9. doi:10.1016/j.devcel.2022.09.002

Barry DM, Xu K, Meadows SM, Zheng Y, Norden PR, Davis GE, Cleaver O. 2015. Cdc42 is required for cytoskeletal support of endothelial cell adhesion during blood vessel formation. Development dev.125260. doi:10.1242/dev.125260

Bentley K, Franco CA, Philippides A, Blanco R, Dierkes M, Gebala V, Stanchi F, Jones M, Aspalter IM, Cagna G, Weström S, Claesson-Welsh L, Vestweber D, Gerhardt H. 2014. The role of differential VE-cadherin dynamics in cell rearrangement during angiogenesis. Nat Cell Biol 16:309–321. doi:10.1038/ncb2926

Bi D, Lopez JH, Schwarz JM, Manning ML. 2015. A density-independent rigidity transition in biological tissues. Nat Phys 11:1074–1079. doi:10.1038/nphys3471

Birdsey GM, Dryden NH, Amsellem V, Gebhardt F, Sahnan K, Haskard DO, Dejana E, Mason JC, Randi AM. 2008. Transcription factor Erg regulates angiogenesis and endothelial apoptosis through VE-cadherin. Blood 111:3498–3506. doi:10.1182/blood-2007-08-105346

Cao J, Ehling M, März S, Seebach J, Tarbashevich K, Sixta T, Pitulescu ME, Werner A-C, Flach B, Montanez E, Raz E, Adams RH, Schnittler H. 2017. Polarized actin and VE-cadherin dynamics regulate junctional remodelling and cell migration during sprouting angiogenesis. Nat Commun 8:2210. doi:10.1038/s41467-017-02373-8

Cao J, Schnittler H. 2019. Putting VE-cadherin into JAIL for junction remodeling. J Cell Sci 132:jcs222893. doi:10.1242/jcs.222893

Carvalho JR, Fortunato IC, Fonseca CG, Pezzarossa A, Barbacena P, Dominguez-Cejudo MA, Vasconcelos FF, Santos NC, Carvalho FA, Franco CA. 2019. Non-canonical Wnt signaling regulates junctional mechanocoupling during angiogenic collective cell migration. eLife 8:e45853. doi:10.7554/eLife.45853

Charlier F, Weber M, Izak D, Harkin E, Magnus M, Lalli J, Fresnais L, Chan M, Markov N, Amsalem O, Proost S, Agamemnon Krasoulis, Getzze, Repplinger S. 2022. trevismd/statannotations: v0.5. doi:10.5281/ZENODO.7213391

Chavkin NW, Genet G, Poulet M, Jeffery ED, Marziano C, Genet N, Vasavada H, Nelson EA, Acharya BR, Kour A, Aragon J, McDonnell SP, Huba M, Sheynkman GM, Walsh K, Hirschi KK. 2022. Endothelial cell cycle state determines propensity for arterial-venous fate. Nat Commun 13:5891. doi:10.1038/s41467-022-33324-7

Chen L. 2009. Ocular lymphatics: state-of-the-art review. Lymphology 42:66–76.

Chrostek A, Wu X, Quondamatteo F, Hu R, Sanecka A, Niemann C, Langbein L, Haase I, Brakebusch C. 2006. Rac1 Is Crucial for Hair Follicle Integrity but Is Not Essential for Maintenance of the Epidermis. Mol Cell Biol 26:6957–6970. doi:10.1128/MCB.00075-06

Claxton S, Fruttiger M. 2005. Oxygen modifies artery differentiation and network morphogenesis in the retinal vasculature. Dev Dyn Off Publ Am Assoc Anat 233:822– 828. doi:10.1002/dvdy.20407

Codling EA, Plank MJ, Benhamou S. 2008. Random walk models in biology. J R Soc Interface 5:813–834. doi:10.1098/rsif.2008.0014

Crawshaw JR, Flegg JA, Bernabeu MO, Osborne JM. 2023. Mathematical models of developmental vascular remodelling: A review. PLOS Comput Biol 19:e1011130. doi:10.1371/journal.pcbi.1011130

Deng Y, Larrivée B, Zhuang ZW, Atri D, Moraes F, Prahst C, Eichmann A, Simons M. 2013. Endothelial RAF1/ERK activation regulates arterial morphogenesis. Blood 121:3988– 3996. doi:10.1182/blood-2012-12-474601

Edgar LT, Park H, Crawshaw JR, Osborne JM, Eichmann A, Bernabeu MO. 2022. Traffic Patterns of the Migrating Endothelium: How Force Transmission Regulates Vascular Malformation and Functional Shunting During Angiogenic Remodelling. Front Cell Dev Biol 10:840066. doi:10.3389/fcell.2022.840066

Eichmann A, Makinen T, Alitalo K. 2005. Neural guidance molecules regulate vascular remodeling and vessel navigation. Genes Dev 19:1013–1021. doi:10.1101/gad.1305405

Ershov D, Phan M-S, Pylvänäinen JW, Rigaud SU, Le Blanc L, Charles-Orszag A, Conway JRW, Laine RF, Roy NH, Bonazzi D, Duménil G, Jacquemet G, Tinevez J-Y. 2022. TrackMate 7: integrating state-of-the-art segmentation algorithms into tracking pipelines. Nat Methods 19:829–832. doi:10.1038/s41592-022-01507-1

Etienne-Manneville S, Hall A. 2002. Rho GTPases in cell biology. Nature 420:629–635. doi:10.1038/nature01148

Fang JS, Coon BG, Gillis N, Chen Z, Qiu J, Chittenden TW, Burt JM, Schwartz MA, Hirschi KK. 2017. Shear-induced Notch-Cx37-p27 axis arrests endothelial cell cycle to enable arterial specification. Nat Commun 8:2149. doi:10.1038/s41467-017-01742-7

Feil R, Brocard J, Mascrez B, LeMeur M, Metzger D, Chambon P. 1996. Ligand-activated site-specific recombination in mice. Proc Natl Acad Sci 93:10887–10890. doi:10.1073/pnas.93.20.10887

Figueiredo AM, Barbacena P, Russo A, Vaccaro S, Ramalho D, Pena A, Lima AP, Ferreira RR, Fidalgo MA, El-Marjou F, Carvalho Y, Vasconcelos FF, Lennon-Duménil A-M, Vignjevic DM, Franco CA. 2021. Endothelial cell invasion is controlled by dactylopodia. Proc Natl Acad Sci 118:e2023829118. doi:10.1073/pnas.2023829118

Franco CA, Jones ML, Bernabeu MO, Geudens I, Mathivet T, Rosa A, Lopes FM, Lima AP, Ragab A, Collins RT, Phng L-K, Coveney PV, Gerhardt H. 2015. Dynamic Endothelial Cell Rearrangements Drive Developmental Vessel Regression. PLOS Biol 13:e1002125. doi:10.1371/journal.pbio.1002125

Gillespie DT. 2007. Stochastic Simulation of Chemical Kinetics. Annu Rev Phys Chem 58:35–55. doi:10.1146/annurev.physchem.58.032806.104637

Gillespie DT. 1977. Exact stochastic simulation of coupled chemical reactions. J Phys Chem 81:2340–2361. doi:10.1021/j100540a008

Giuggioli L, Potts JR, Rubenstein DI, Levin SA. 2013. Stigmergy, collective actions, and animal social spacing. Proc Natl Acad Sci 110:16904–16909. doi:10.1073/pnas.1307071110

Grebenkov DS, Metzler R, Oshanin G. 2019. Full distribution of first exit times in the narrow escape problem. New J Phys 21:122001. doi:10.1088/1367-2630/ab5de4

Heynen SR, Meneau I, Caprara C, Samardzija M, Imsand C, Levine EM, Grimm C. 2013. CDC42 Is Required for Tissue Lamination and Cell Survival in the Mouse Retina. PLoS ONE 8:e53806. doi:10.1371/journal.pone.0053806

Holcman D, Schuss Z. 2014. The Narrow Escape Problem. SIAM Rev 56:213–257. doi:10.1137/120898395

Ivins S, Chappell J, Vernay B, Suntharalingham J, Martineau A, Mohun TJ, Scambler PJ. 2015. The CXCL12/CXCR4 Axis Plays a Critical Role in Coronary Artery Development. Dev Cell 33:455–468. doi:10.1016/j.devcel.2015.03.026

Jin Y, Ding Y, Richards M, Kaakinen M, Giese W, Baumann E, Szymborska A, Rosa A, Nordling S, Schimmel L, Akmeriç EB, Pena A, Nwadozi E, Jamalpour M, Holstein K, Sáinz-Jaspeado M, Bernabeu MO, Welsh M, Gordon E, Franco CA, Vestweber D, Eklund L, Gerhardt H, Claesson-Welsh L. 2022. Tyrosine-protein kinase Yes controls endothelial junctional plasticity and barrier integrity by regulating VE-cadherin phosphorylation and endocytosis. Nat Cardiovasc Res. doi:10.1038/s44161-022-00172-z

Karaiskos N, Wahle P, Alles J, Boltengagen A, Ayoub S, Kipar C, Kocks C, Rajewsky N, Zinzen RP. 2017. The *Drosophila* embryo at single-cell transcriptome resolution. Science 358:194–199. doi:10.1126/science.aan3235

Kim YH, Hu H, Guevara-Gallardo S, Lam MTY, Fong S-Y, Wang RA. 2008. Artery and vein size is balanced by Notch and ephrin B2/EphB4 during angiogenesis. Development 135:3755–3764. doi:10.1242/dev.022475

Laviña B, Castro M, Niaudet C, Cruys B, Álvarez-Aznar A, Carmeliet P, Bentley K, Brakebusch C, Betsholtz C, Gaengel K. 2018. Defective endothelial cell migration in the absence of Cdc42 leads to capillary-venous malformations. Development dev.161182. doi:10.1242/dev.161182

Lawson CD, Ridley AJ. 2018. Rho GTPase signaling complexes in cell migration and invasion. J Cell Biol 217:447–457. doi:10.1083/jcb.201612069

Lee H-W, Shin JH, Simons M. 2022. Flow goes forward and cells step backward: endothelial migration. Exp Mol Med 54:711–719. doi:10.1038/s12276-022-00785-1

Lee H-W, Xu Y, He L, Choi W, Gonzalez D, Jin S-W, Simons M. 2021. Role of Venous Endothelial Cells in Developmental and Pathologic Angiogenesis. Circulation 144:1308–1322. doi:10.1161/CIRCULATIONAHA.121.054071

Lenne P-F, Trivedi V. 2022. Sculpting tissues by phase transitions. Nat Commun 13:664. doi:10.1038/s41467-022-28151-9

Liang J, Oyang L, Rao S, Han Y, Luo X, Yi P, Lin J, Xia L, Hu J, Tan S, Tang L, Pan Q, Tang Y, Zhou Y, Liao Q. 2021. Rac1, A Potential Target for Tumor Therapy. Front Oncol 11:674426. doi:10.3389/fonc.2021.674426

Luo W, Garcia-Gonzalez I, Fernández-Chacón M, Casquero-Garcia V, Sanchez-Muñoz MS, Mühleder S, Garcia-Ortega L, Andrade J, Potente M, Benedito R. 2021. Arterialization requires the timely suppression of cell growth. Nature 589:437–441. doi:10.1038/s41586-020-3018-x

Martinez-Corral I, Stanczuk L, Frye M, Ulvmar MH, Diéguez-Hurtado R, Olmeda D, Makinen T, Ortega S. 2016. Vegfr3-CreER T2 mouse, a new genetic tool for targeting the lymphatic system. Angiogenesis 19:433–445. doi:10.1007/s10456-016-9505-x

Maseda F, Cang Z, Nie Q. 2021. DEEPsc: A Deep Learning-Based Map Connecting Single-Cell Transcriptomics and Spatial Imaging Data. Front Genet 12:636743. doi:10.3389/fgene.2021.636743

Mirams GR, Pathmanathan P, Gray RA, Challenor P, Clayton RH. 2016. Uncertainty and variability in computational and mathematical models of cardiac physiology: Uncertainty and variability in cardiac models. J Physiol 594:6833–6847. doi:10.1113/JP271671

Montroll EW, Weiss GH. 1965. Random Walks on Lattices. II. J Math Phys 6:167–181. doi:10.1063/1.1704269

Mukouyama Y, Shin D, Britsch S, Taniguchi M, Anderson DJ. 2002. Sensory Nerves Determine the Pattern of Arterial Differentiation and Blood Vessel Branching in the Skin. Cell 109:693–705. doi:10.1016/S0092-8674(02)00757-2

Muzumdar MD, Tasic B, Miyamichi K, Li L, Luo L. 2007. A global double-fluorescent Cre reporter mouse. genesis 45:593–605. doi:10.1002/dvg.20335

Neto F, Klaus-Bergmann A, Ong YT, Alt S, Vion A-C, Szymborska A, Carvalho JR, Hollfinger I, Bartels-Klein E, Franco CA, Potente M, Gerhardt H. 2018. YAP and TAZ regulate adherens junction dynamics and endothelial cell distribution during vascular development. eLife 7:e31037. doi:10.7554/eLife.31037

Paatero I, Sauteur L, Lee M, Lagendijk AK, Heutschi D, Wiesner C, Guzmán C, Bieli D, Hogan BM, Affolter M, Belting H-G. 2018. Junction-based lamellipodia drive endothelial cell rearrangements in vivo via a VE-cadherin-F-actin based oscillatory cell-cell interaction. Nat Commun 9:3545. doi:10.1038/s41467-018-05851-9

Pedregosa F, Varoquaux G, Gramfort A, Michel V, Thirion B, Grisel O, Blondel M, Prettenhofer P, Weiss R, Dubourg V, others. 2011. Scikit-learn: Machine learning in Python. J Mach Learn Res 12:2825–2830.

Phng L-K, Gebala V, Bentley K, Philippides A, Wacker A, Mathivet T, Sauteur L, Stanchi F, Belting H-G, Affolter M, Gerhardt H. 2015. Formin-Mediated Actin Polymerization at Endothelial Junctions Is Required for Vessel Lumen Formation and Stabilization. Dev Cell 32:123–132. doi:10.1016/j.devcel.2014.11.017

Phng L-K, Stanchi F, Gerhardt H. 2013. Filopodia are dispensable for endothelial tip cell guidance. Development 140:4031–4040. doi:10.1242/dev.097352

Pitulescu ME, Schmidt I, Giaimo BD, Antoine T, Berkenfeld F, Ferrante F, Park H, Ehling M, Biljes D, Rocha SF, Langen UH, Stehling M, Nagasawa T, Ferrara N, Borggrefe T, Adams RH. 2017. Dll4 and Notch signalling couples sprouting angiogenesis and artery formation. Nat Cell Biol 19:915–927. doi:10.1038/ncb3555

Raftopoulou M, Hall A. 2004. Cell migration: Rho GTPases lead the way. Dev Biol 265:23–32. doi:10.1016/j.ydbio.2003.06.003

Ridley AJ. 2001. Rho family proteins: coordinating cell responses. Trends Cell Biol 11:471–477. doi:10.1016/S0962-8924(01)02153-5

Rosa A, Giese W, Meier K, Alt S, Klaus-Bergmann A, Edgar LT, Bartels-Klein E, Collins RT, Szymborska A, Coxam B, Bernabeu MO, Gerhardt H. 2022. WASp controls oriented migration of endothelial cells to achieve functional vascular patterning. Development 149:dev200195. doi:10.1242/dev.200195

Rust R, Grönnert L, Dogançay B, Schwab ME. 2019. A Revised View on Growth and Remodeling in the Retinal Vasculature. Sci Rep 9:3263. doi:10.1038/s41598-019-40135-2

Satija R, Farrell JA, Gennert D, Schier AF, Regev A. 2015. Spatial reconstruction of single-cell gene expression data. Nat Biotechnol 33:495–502. doi:10.1038/nbt.3192

Sauteur L, Krudewig A, Herwig L, Ehrenfeuchter N, Lenard A, Affolter M, Belting H-G. 2014. Cdh5/VE-cadherin Promotes Endothelial Cell Interface Elongation via Cortical Actin Polymerization during Angiogenic Sprouting. Cell Rep 9:504–513. doi:10.1016/j.celrep.2014.09.024

Schindelin J, Arganda-Carreras I, Frise E, Kaynig V, Longair M, Pietzsch T, Preibisch S, Rueden C, Saalfeld S, Schmid B, Tinevez J-Y, White DJ, Hartenstein V, Eliceiri K, Tomancak P, Cardona A. 2012. Fiji: an open-source platform for biological-image analysis. Nat Methods 9:676–682. doi:10.1038/nmeth.2019

Schmidt U, Weigert M, Broaddus C, Myers G. 2018. Cell Detection with Star-Convex Polygons In: Frangi AF, Schnabel JA, Davatzikos C, Alberola-López C, Fichtinger G, editors. Medical Image Computing and Computer Assisted Intervention – MICCAI 2018, Lecture Notes in Computer Science. Cham: Springer International Publishing. pp. 265–273. doi:10.1007/978-3-030-00934-2_30

Scott DW. 1979. On optimal and data-based histograms. Biometrika 66:605–610. doi:10.1093/biomet/66.3.605

Stepanova D, Byrne HM, Maini PK, Alarcón T. 2021. A multiscale model of complex endothelial cell dynamics in early angiogenesis. PLOS Comput Biol 17:e1008055. doi:10.1371/journal.pcbi.1008055

Su T, Stanley G, Sinha R, D’Amato G, Das S, Rhee S, Chang AH, Poduri A, Raftrey B, Dinh TT, Roper WA, Li G, Quinn KE, Caron KM, Wu S, Miquerol L, Butcher EC, Weissman I, Quake S, Red-Horse K. 2018. Single-cell analysis of early progenitor cells that build coronary arteries. Nature 559:356–362. doi:10.1038/s41586-018-0288-7

Tabibian A, Ghaffari S, Vargas DA, Van Oosterwyck H, Jones EAV. 2020. Simulating flow induced migration in vascular remodelling. PLOS Comput Biol 16:e1007874. doi:10.1371/journal.pcbi.1007874

Tammela T, Alitalo K. 2010. Lymphangiogenesis: Molecular Mechanisms and Future Promise. Cell 140:460–476. doi:10.1016/j.cell.2010.01.045

Tammela T, Zarkada G, Nurmi H, Jakobsson L, Heinolainen K, Tvorogov D, Zheng W, Franco CA, Murtomäki A, Aranda E, Miura N, Ylä-Herttuala S, Fruttiger M, Mäkinen T, Eichmann A, Pollard JW, Gerhardt H, Alitalo K. 2011. VEGFR-3 controls tip to stalk conversion at vessel fusion sites by reinforcing Notch signalling. Nat Cell Biol 13:1202–1213. doi:10.1038/ncb2331

Tan W, Palmby TR, Gavard J, Amornphimoltham P, Zheng Y, Gutkind JS. 2008. An essential role for Rac1 in endothelial cell function and vascular development. FASEB J Off Publ Fed Am Soc Exp Biol 22:1829–1838. doi:10.1096/fj.07-096438

Thompson JF. 1982. General curvilinear coordinate systems. Appl Math Comput 10–11:1–30. doi:10.1016/0096-3003(82)90185-0

Trepat X, Sahai E. 2018. Mesoscale physical principles of collective cell organization. Nat Phys 14:671–682. doi:10.1038/s41567-018-0194-9

Tzima E. 2002. Activation of Rac1 by shear stress in endothelial cells mediates both cytoskeletal reorganization and effects on gene expression. EMBO J 21:6791–6800. doi:10.1093/emboj/cdf688

Tzima E, Kiosses WB, Del Pozo MA, Schwartz MA. 2003. Localized Cdc42 Activation, Detected Using a Novel Assay, Mediates Microtubule Organizing Center Positioning in Endothelial Cells in Response to Fluid Shear Stress. J Biol Chem 278:31020– 31023. doi:10.1074/jbc.M301179200

Ueda HR, Dodt H-U, Osten P, Economo MN, Chandrashekar J, Keller PJ. 2020. Whole-Brain Profiling of Cells and Circuits in Mammals by Tissue Clearing and Light-Sheet Microscopy. Neuron 106:369–387. doi:10.1016/j.neuron.2020.03.004

Vion A-C, Perovic T, Petit C, Hollfinger I, Bartels-Klein E, Frampton E, Gordon E, Claesson-Welsh L, Gerhardt H. 2021. Endothelial Cell Orientation and Polarity Are Controlled by Shear Stress and VEGF Through Distinct Signaling Pathways. Front Physiol 11:623769. doi:10.3389/fphys.2020.623769

Weijts B, Gutierrez E, Saikin SK, Ablooglu AJ, Traver D, Groisman A, Tkachenko E. 2018. Blood flow-induced Notch activation and endothelial migration enable vascular remodeling in zebrafish embryos. Nat Commun 9:5314. doi:10.1038/s41467-018-07732-7

Wu X, Quondamatteo F, Lefever T, Czuchra A, Meyer H, Chrostek A, Paus R, Langbein L, Brakebusch C. 2006. Cdc42 controls progenitor cell differentiation and β-catenin turnover in skin. Genes Dev 20:571–585. doi:10.1101/gad.361406

Xu C, Hasan SS, Schmidt I, Rocha SF, Pitulescu ME, Bussmann J, Meyen D, Raz E, Adams RH, Siekmann AF. 2014. Arteries are formed by vein-derived endothelial tip cells. Nat Commun 5:5758. doi:10.1038/ncomms6758

Zegers MM, Friedl P. 2014. Rho GTPases in collective cell migration. Small GTPases 5:e983869. doi:10.4161/sgtp.28997

