## Supplemental Text S1 for "Reconstructing stochastic cell population trajectories reveals regulators and heterogeneity of endothelial flow-migration coupling driving vascular remodelling"

### Supplementary Text S1: Reconstructing stochastic cell population trajectories reveals regulators and heterogeneity of endothelial flow-migration coupling driving vascular remodelling

#### Contents

**1 Model description** **1**
1.1 Distance notations and domains . . . . . . . . . . . . . . . . . . . . . . . . . . 2
1.2 Force field computation from image data . . . . . . . . . . . . . . . . . . . . . 3

**2 Bibliography** **4**

**3 Parameters** **5**

### 1 Model description

The movement of endothelial cell (EC) agents from one lattice site to another is described by a step selection function. The step selection function $f_i^{\tau,t}(\vec{x}|\vec{y})$ describes the probability of an EC agent $i$ moving from position $\vec{x}$ at time $t$ to position $\vec{y}$ at time $t+\tau$. In the normal case we will consider $\vec{x}=(x_1,x_2)$ and $\vec{y}=(y_1,y_2)$ as two-dimensional vectors. The step selection function comprises three different aspects of cellular movement, which are (i) biased Brownian movement , (ii) signalling cues, (iii) mechanical cues and the (iv) cellular state. In this model we will neglect cell-cell interactions.

The model simulations are based on the spatial Gillespie algorithm [1], which was originally used for reaction-diffusion systems in molecular dynamics. The model framework has recently been adapted to cellular and animal movement [2, 3]. We will describe cellular movement in terms of rates which express the expected time of an EC agent moving from one lattice site to another. Given that an EC agent $i$ is at site $\vec{x}$, the probability distribution for an agent moving from site $\vec{x}$. to site $\vec{y}$ is given by:

$$f_i^{\tau,t}(\vec{x}|\vec{y},t) \propto \mu_{i,k}(\vec{x}|\vec{y}) \exp(-\tau\mu_i),$$

The corresponding rates of movement are given by

$$\mu_{i,k}(\vec{x}|\vec{y},t) = \begin{cases} \frac{\mu_i}{4}\left[1+\alpha\frac{(\vec{x}-\vec{y})}{|x-y|}\cdot\vec{b}(\vec{x})\right], & \text{if } |\vec{x}-\vec{y}|=a, \\ 0 & \text{else.} \end{cases} \tag{1}$$

Here, $\vec{b}(\vec{x})$ determines the bias of an EC agent to move in direction of $\vec{b}(\vec{x})$. In case of $b(\vec{x}) \equiv 0$, the EC agent becomes a random walker with diffusion coefficient $D_i = h^2 \frac{\mu_i}{4}$ and the instantaneous velocity is given by $v = \mu_i h$, where $h$ is the grid size.

#### 1.1 Distance notations and domains

The simulation is performed on a vasculature from retinal image data. We introduce the following distance notations:

$$d_a(\vec{x}) = \inf_{y \in \Omega_A} d(\vec{x}, y), \tag{2}$$

$$d_v(\vec{x}) = \inf_{y \in \Omega_V} d(\vec{x}, y), \tag{3}$$

$$r(\vec{x}) = \inf_{y \in \Omega_{ON}} d(\vec{x}, y). \tag{4}$$

Here, $\Omega_A, \Omega_V$ and $\Omega_{ON}$ are the labelled domains of artery, vein and optic nerve. The normalized vein-artery distance $\phi_{v-a,rp}$ is computed as follows

$$\phi_{v-a,rp}(\vec{x}) = \frac{d_v(\vec{x})}{d_a(\vec{x}) + d_v(\vec{x})}, \tag{5}$$

The transition of remodelling plexus to sprouting front is described by the radial distance $r_{plex}(t)$, which is a function of time

$$r_{rp}(t) = r_{rp}^0 + v_{rp} t. \tag{6}$$

#### 1.2 Force field computation from image data

Using these distance notations, the shear stress cue is computed from

$$\vec{b}_{ss} = \vec{a}_{ss} + \vec{v}_{ss} + \nabla U_{ss}, \tag{7}$$

$$U_{ss} = \phi_{v-a,rp} + \frac{1}{4}\phi^2_{v-a,rp}, \tag{8}$$

$$\vec{a}_{ss} = \frac{f_a}{k_a + d_a}\nabla r, \tag{9}$$

$$\vec{v}_{ss} = -\frac{f_v}{k_v + d_v}\nabla r. \tag{10}$$

The directed VEGF-A cue is computed from the distance to the artery mask in the following way:

$$b_{VEGF} = \nabla \left(d_a^2(x)\right) + \rho \nabla r. \tag{11}$$

The shear stress cue can overlap into the sprouting front and the VEGFA cue into the remodelling plexus, which defines a competition region between both forces [4]. We define the functions

$$g_{ss}(\vec{x}) = 1 - \frac{r^\gamma(\vec{x})}{(r_{rp} + \rho_{ss})^\gamma + r^\gamma(\vec{x})}, \tag{12}$$

$$g_{VEGFA}(\vec{x}) = \frac{r^\gamma(\vec{x})}{(r_{rp} - \rho_{\text{VEGFA}})^\gamma + r^\gamma(\vec{x})}. \tag{13}$$

The resulting shear stress and VEGFA cues are then scaled in the following way

$$\vec{b}^E_{ss}(\vec{x}) = g_{ss}(\vec{x})\frac{\vec{b}_{ss}(\vec{x})}{|\vec{b}_{ss}(\vec{x})|} \tag{14}$$

and

$$\vec{b}^E_{VEGFA}(\vec{x}) = g_{VEGFA}(\vec{x})\frac{\vec{b}_{VEGFA}(\vec{x})}{|\vec{b}_{VEGFA}(\vec{x})|}. \tag{15}$$

The resulting cue is then computed from

$$\vec{b}(\vec{x}) = \alpha[\xi \vec{b}^E_{ss}(\vec{x}) + (1-\xi)\vec{b}^E_{VEGFA}(\vec{x})]. \tag{16}$$

### 2 Bibliography

#### References


[1] Sotiria Lampoudi, Dan T Gillespie, and Linda R Petzold. The multinomial simulation algorithm for discrete stochastic simulation of reaction-diffusion systems. *The Journal of chemical physics*, 130(9):094104, 2009.

[2] Daria Stepanova, Helen M Byrne, Philip K Maini, and Tomás Alarcón. A multiscale model of complex endothelial cell dynamics in early angiogenesis. *PLoS Computational Biology*, 17(1):e1008055, 2021.

[3] Luca Giuggioli, Jonathan R Potts, Daniel I Rubenstein, and Simon A Levin. Stigmergy, collective actions, and animal social spacing. *Proceedings of the National Academy of Sciences*, 110(42):16904–16909, 2013.

[4] Pedro Barbacena, Maria Dominguez-Cejudo, Catarina G Fonseca, Manuel Gómez-González, Laura M Faure, Georgia Zarkada, Andreia Pena, Anna Pezzarossa, Daniela Ramalho, Ylenia Giarratano, et al. Competition for endothelial cell polarity drives vascular morphogenesis in the mouse retina. *Developmental Cell*, 57(19):2321–2333, 2022.

### 3 Parameters

| parameter | Value | description |
|---|---|---|
| $D$ | - | diffusion coefficient |
| $\alpha$ | - | strength of coupling to dual force field |
| $\xi$ | - | fraction of shear stress cue |
| $\lambda_{ss}$ | 75 $\mu$m | overlap of shear stress into sprouting front |
| $\lambda_{\text{VEGFA}}$ | 125 $\mu$m | overlap of shear stress into sprouting front |
| $a$ | 0.69 $\mu$m | size of one grid element |
| $f_a$ | 0.05 | relative strength of shear stress cue in artery |
| $k_a$ | 10 $\mu$m | - |
| $f_v$ | 0.05 | relative strength of shear stress cue in vein |
| $k_v$ | 10 $\mu$m | - |
| $r^0_{plex}$ | 900 $\mu$m | initial end of the remodelling plexus and start of sprouting front |
| $v_{plex}$ | 5 $\mu m/h$ | speed of sprouting front progression |
| $\gamma$ | 10 | parameter for steepness of competition zone |

Table 1: Parameters used in the model.

| parameter | Model-Ctr | Model-Cdc42 | Model-Rac1 |
|---|---|---|---|
| $\mu_r$ | 18 steps per h | 18 steps per h | 18 steps per h |
| $\alpha$ | 0.36 | 0.26 | 0.32 |
| $\xi$ | 0.53 | 0.19 | 0.34 |

Table 2: Optimized parameters.
